# Cell lineage analysis with somatic mutations reveals late divergence of neuronal cell types and cortical areas in human cerebral cortex

**DOI:** 10.1101/2023.11.06.565899

**Authors:** Sonia Nan Kim, Vinayak V. Viswanadham, Ryan N. Doan, Yanmei Dou, Sara Bizzotto, Sattar Khoshkhoo, August Yue Huang, Rebecca Yeh, Brian Chhouk, Alex Truong, Kathleen M. Chappell, Marc Beaudin, Alison Barton, Shyam K. Akula, Lariza Rento, Michael Lodato, Javier Ganz, Ryan A. Szeto, Pengpeng Li, Jessica W. Tsai, Robert Sean Hill, Peter J. Park, Christopher A. Walsh

## Abstract

The mammalian cerebral cortex shows functional specialization into regions with distinct neuronal compositions, most strikingly in the human brain, but little is known in about how cellular lineages shape cortical regional variation and neuronal cell types during development. Here, we use somatic single nucleotide variants (sSNVs) to map lineages of neuronal sub-types and cortical regions. Early-occurring sSNVs rarely respect Brodmann area (BA) borders, while late-occurring sSNVs mark neuron-generating clones with modest regional restriction, though descendants often dispersed into neighboring BAs. Nevertheless, in visual cortex, BA17 contains 30-70% more sSNVs compared to the neighboring BA18, with clones across the BA17/18 border distributed asymmetrically and thus displaying different cortex-wide dispersion patterns. Moreover, we find that excitatory neuron-generating clones with modest regional restriction consistently share low-mosaic sSNVs with some inhibitory neurons, suggesting significant co-generation of excitatory and some inhibitory neurons in the dorsal cortex. Our analysis reveals human-specific cortical cell lineage patterns, with both regional inhomogeneities in progenitor proliferation and late divergence of excitatory/inhibitory lineages.

## Introduction

Pattern formation in the human cerebral cortex has fascinated neuroscientists for more than a century. Cortical areas, which were originally defined by cytoarchitectonic and myeloarchitectonic differences encapsulated in Brodmann Area (BAs) designations (Brodmann, 1909) over a century ago, show dramatic changes in relative cellular components across functional boundaries in human cortex (Brodmann, 1909; Budday et al., 2015). Recent investigations of large human cohorts with MRI measures of function and connectivity (Glasser et al., 2016) have largely confirmed the existence of dozens of functional specializations that correlate with BAs, identified dozens more, and validated their relevance for complex cognitive and behavioral tasks (Binkofski and Buccino, 2004; Hanakawa et al., 2002). Diverse cortical areas appear to arise by a complex interplay of patterning forces, as exemplified by the primate visual cortex. Early regional specification of cortex relies on gradients of secreted factors and transcription factors (Ypsilanti and Rubenstein, 2016), without explicit requirement for axonal input (Cadwell et al., 2019). However, later differentiation of specific regions shows roles for extrinsic cues. For example, prenatal removal of visual thalamic inputs in monkeys causes a reduction in the extent of the primary visual cortex (BA17), along with a shift of the histologically identifiable boundary between the primary visual cortex and secondary visual cortex (BA18) (Dehay et al., 1996, 1989), suggesting that extrinsic factors can also regulate cortical area specification, cell identity, and connectivity (Dehay et al., 1989). Thus, both intrinsic and extrinsic variables help specify the cytoarchitecture and cell types in the primate visual system (Dehay et al., 1996, 1989; Rakic et al., 1991). Direct studies of cell lineage in animal models have given variable results about potential roles of lineage in regional and cell-type specification. In the mouse, radial glia (which produce excitatory neurons of the cortex) have been proposed to produce neurons that are preferentially connected to each other, distributed in both the superficial and deep layers of the mouse cortex (Rakic, 1988; Gao et al., 2014) and sharing some functional properties (Rakic et al., 1991; Gao et al., 2014; Li et al., 2012; Ohtsuki et al., 2012). Such an observation may suggest that cellular lineage contributes to specifying cortical circuits. In contrast, other studies have suggested stochasticity in mouse cortical neurogenesis, with pyramidal neuron clones showing a wide range of sizes and laminar configurations, including deep layer-restricted cortical lineages (Yu et al., 2009). However, the extent to which lineages in different cortical areas may exhibit features relating to the local cytoarchitecture and leading to heterogeneous progeny is unknown.

To address the questions on the spatial outcomes and identities of cells produced by human cerebral cortical progenitor cells, we utilized somatic mutations as endogenous markers of cellular relatedness. Lineage and clonal analyses using somatic single-nucleotide variants (sSNVs) identified in postmortem human brain samples have shed light into the earliest events of brain development and cell type relationships in the prefrontal cortex, all pointing to a large degree of clonal dispersion (Bizzotto et al., 2021; Lodato et al., 2015; Huang et al., 2020; Breuss et al., 2022). Here, to describe human-specific cortical lineage patterns, we characterized the spatial distribution of cellular clones across the human cerebral cortex using sSNVs and integrated this data with transcriptomic information about cells in each clone. We confirmed earlier reports of widespread clonal dispersion across human cortex, with even late-rising mosaic variants present across multiple regions. However, we also discovered regional inhomogeneity in clonal structure superimposed on this dispersion, notably at the BA17/18. Surprisingly, we also observed the frequent co-generation of excitatory and inhibitory neurons at late stages of neurogenesis (Delgado et al., 2021), providing the first demonstration that these mixed clones are common in vivo.

## Results

### The primary visual cortex harbors 30-70% more sSNVs than the adjacent secondary visual cortex

We first sought to identify and validate clonal mosaic sSNVs in the cortex before tracking them across multiple brain regions and using them to reconstruct cellular lineages (**Fig. 1A**). As described before (Bizzotto et al., 2021; Dou et al., 2020), we discovered sSNVs by applying MosaicForecast (a machine learning algorithm we developed (Dou et al., 2020)) to very high-coverage (210X) whole-genome sequencing (WGS) data of bulk DNA extracted from four neurotypical individuals. In each individual, we profiled grey matter of three regions: BA9 within the prefrontal cortex (PFC) and two areas from the occipital lobe (primary visual cortex, BA17, and secondary visual cortex, BA18; **Fig. 1B**; **Table S1.1**; **Methods**).

**Figure 1:**
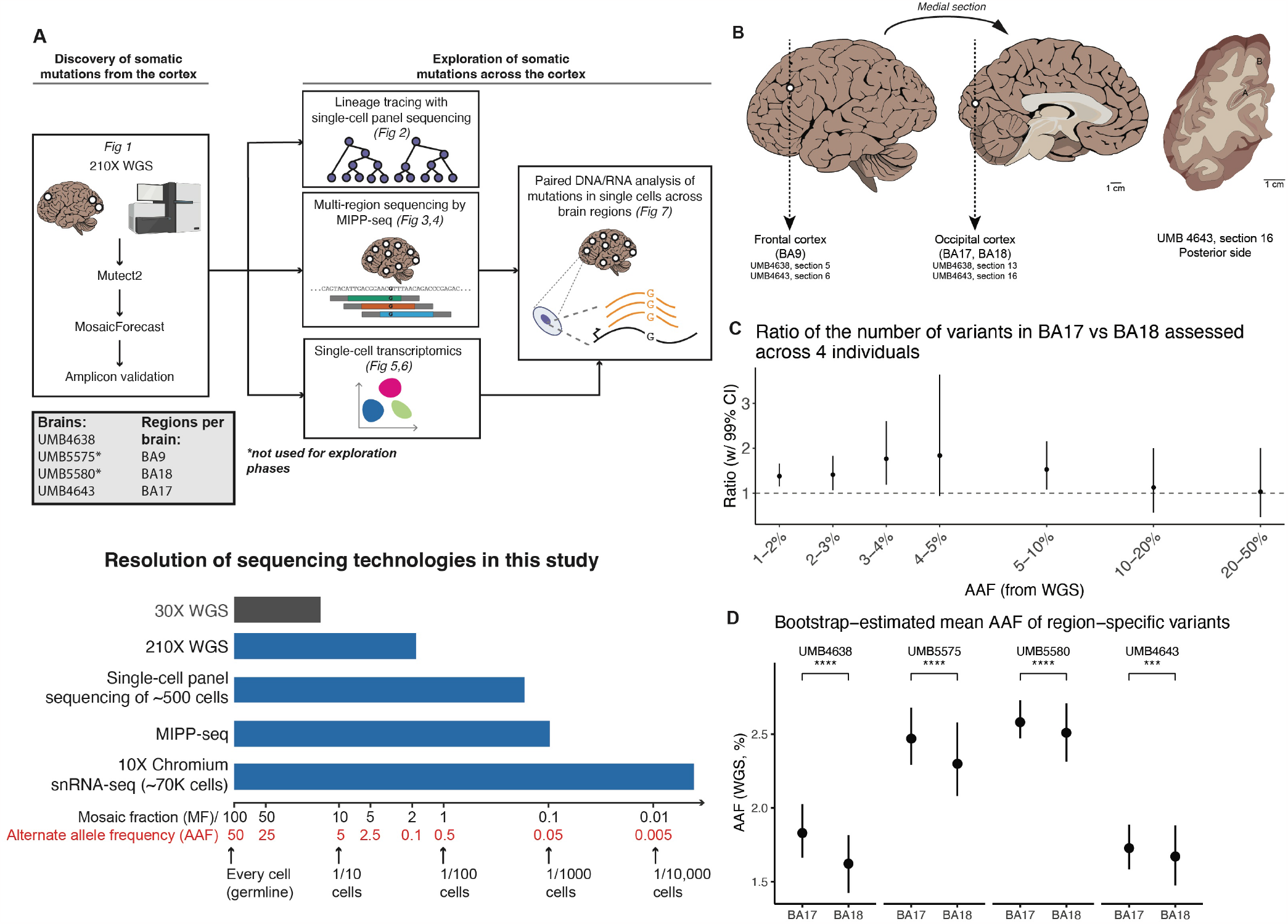
Regional differences in sSNV counts between cortical areas. **(A)** *Top*: Experimental outline for clonal sSNV analysis, with each analysis corresponding to a relevant figure in this paper. *Bottom*: A schematic comparison of the resolution of different sequencing technologies used in this study (blue) for variants present at different mosaic fractions (and corresponding alternate allele fraction) in tissue. 30X WGS (not used for this study) is shown in grey for comparison. **(B)** DNA was prepared and sequenced from the frontal (BA9) and occipital cortex (BA18 and BA17). Tracings of the occipital cortex section show sampling of adjacent cortical areas: primary visual cortex, BA17 (A) indicated by the stria of Gennari (white line); and secondary visual cortex, BA18 (B). **(C)** Ratio of the number of variants between the adjacent regions (BA18 and BA17) per alternative allele frequency (AAF) range. Calculations were performed across all 4 brains (UMB4638, UMB4643, UMB5575, and UMB5580). The “control” distributions were generated from the ratios of the simulated numbers of mutations drawn from each region if the corresponding variants were present at the average of the AAFs at which they were discovered in BA17 and BA18. **(D)** Bootstrap estimates of the AAFs of variants specific to either BA17 or BA18. The significance of AAF estimates is assessed by the t-test (***: p ¡ 1e-3; ****: p ¡ 1e-4).

Across all four individuals, we found that BA17 consistently showed more sSNV than the adjacent BA18, with BA9 also exceeding BA18 in 3/4 samples (**Fig. 1C,D**; **Fig. S1A-D**). We focused on the differences between adjoining BA17 and BA18 first, estimating the ratio of sSNVs detectable in BA17 versus BA18 at different alternate allele fractions (AAF) after controlling for the sensitivity to detect sSNVs in deep bulk WGS data (**Methods**). We found that BA17 contains 30-70% more sSNVs than BA18 at <5% AAF (equivalently, at mosaic fractions (MFs) of <10%; **Fig. 1A**), suggesting that the excess of sSNVs in BA17 over BA18 arises near the start of and persists throughout cortical neurogenesis. In contrast, sSNVs at AAFs >10%, likely arising in early embryonic cell divisions before the formation of brain16 were distributed equally in all three regions (**Fig. 1C**; **Fig. S1C**). The greater number of sSNVs in BA9 over BA18 in three of the brains (UMB4638, UMB4643, and UMB5575; **Fig. S1A-C**), may be reflecting the frontal cortex’s large size, distance from the occipital cortex, and relatively late generation, all of which would lead to larger sSNV counts (Kolk and Rakic, 2022). In addition to the consistently greater number of sSNVs detected in BA17 versus BA18, we found that BA17-restricted variants were present at higher average AAFs than BA18-restricted variants (**Fig. 1D**). Variants shared between the two regions did not have significantly different average AAFs in one region versus the other (**Fig. S1D**). Region-restricted variants were detected at AAFs of ≈ 1.5-2.5% and shared-region variants at≈ 6-15%, consistent with region-specific variants arising later than shared variants.

Amplicon panel sequencing (>10,000X; **Methods**) confirmed the >1.5x higher number of sSNVs in BA17 compared to BA18. We selected a set of 155 total sSNVs originally detected in BA17 and/or BA18 (**Table S1.1-1.5**). Of the 155 sSNVs, 138 (89%) were validated in the original discovery site (ODS) (**Table S1.2**). More than half of the variants at <3% AAF (<6% MF) were specific to the brain (**Fig. S1E**; **Table S1.3**), in line with previously-published analyses showing that most brain-restricted SNVs appear at AAFs of <1% with some appearing at 1-5% AAF (Bizzotto et al., 2021). Our overall sSNV validation rate was as expected from MosaicForecast-based discovery (Dou et al., 2020; Rodin et al., 2021) (**Table S1.3-1.5**).

The two consistent trends—BA17 containing more sSNVs than BA18, and BA17-restricted variants occurring at higher AAFs than BA18-restricted variants—suggest a fundamental difference in the clonal structures of these two neighboring regions. Assuming that our sSNVs are neutral to clonal selection, the higher number of sSNV in BA17 could reflect a larger progenitor pool than in BA18, and would presumably also require that some progeny remain mostly regionally restricted. The greater AAFs of BA17-restricted variants could arise from BA17 gaining its region-restricted progenitors earlier than BA18 and thus having more time to clonally expand and show sSNVs at higher cell fractions. These patterns are consistent with previous observations in non-human primates of a higher density of neurons in adult primary compared to secondary visual cortex (Collins et al., 2010; Dehay and Kennedy, 2007; Rockel et al., 1980) and constrained radial migration across the border during development (Li et al., 2012; Ohtsuki et al., 2012; Yu et al., 2009). Studies of non-human primates also suggest that the proliferation rate is greater for progenitors underlying the incipient primary visual cortex compared to the secondary visual cortex (Dehay et al., 1993; Smart et al., 2002; Lukaszewicz et al., 2005).

### Inferring the timing of cortical patterning using single-cell lineage tracing

To infer a more detailed timeline of cortical spatial patterning, we generated lineage trees from 122 brain-restricted SNVs profiled by panel sequencing in 1131 single cells taken from BA17, BA18, and BA9 (as an outgroup) (**Methods**; **Fig. S2AD**). To estimate the time for all observed cells to develop from their most recent common ancestor (MRCA), we computed the coalescent time of the population (**Methods**; **Supplemental Methods**; **Fig. S2E**). Each variant arises at some point during the population’s development, and the cell in which the variant first arises will yield progeny forming a subpopulation in our observed cells. Thus, the coalescent time of a subpopulation can be used to estimate the time-of-origin (TOO) of its corresponding variant. Using published values for the number of new cells generated during human fetal neurogenesis (Ackerman, 1992), we converted each variant’s coalescent time to an estimate of chronological time. We present each variant’s TOO as the number of weeks that have elapsed since the MRCA of the entire corresponding lineage (“post-MRCA week,” or PMW).

We estimated that analyzed neurons in each brain formed their lineages over 17 weeks (14.8 weeks for UMB4638 and 18.7 weeks for UMB4643; **Fig. S2D**). This timespan corresponds roughly to the length of human cortical neurogenesis (gestational weeks [GW] 10-25) (Stepien et al., 2021; Malik et al., 2013; Sidman and Rakic, 1973). If our MRCAs first arose at or near the start of neurogenesis (i.e., PMW0 = GW10), then our lineages would track developmental events occurring throughout most of neurogenesis. Our cells were sampled from 54-68 million cells within which the mutations studied in each tree may be found, with 7 new mutations arising in the population during each generation (**Fig. S2E**). Our lineage analysis provides estimated TOOs for variants, the order of variants along successive branches in the lineage tree, and the observed regional distributions of cells carrying a subpopulation-defining variant. With this information, we constructed timelines of variants that track how descendants of cortical clones spread out across the prefrontal and occipital cortex (**Fig. 2A**; **Methods**).

**Figure 2:**
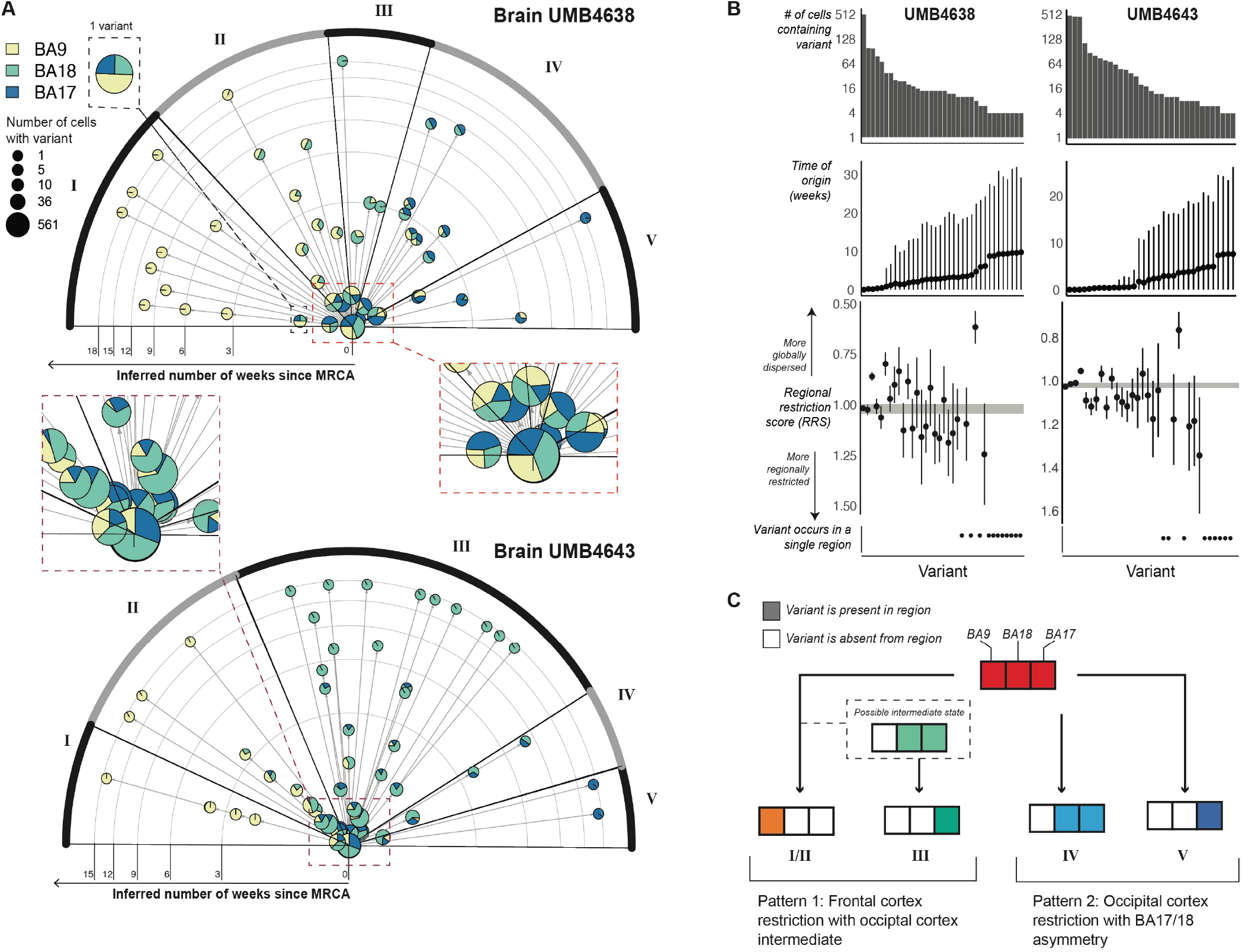
Timelines of variant occurrence across the visual and prefrontal cortices as estimated from single-cell lineage trees and coalescent models. **(A)** Radial plot showing the time-of-origin (TOO) of variants inferred by a coalescent model applied to single-cell lineages. For each brain, a single-cell lineage tree was built on 580 cells and 40-50 variants. Each variant is represented by a pie chart sized by the number of cells carrying the variant and sliced by the breakdown of regions where these cells were found. If one variant occurs right before another within the same lineage (i.e., on consecutive branches of the lineage tree), then an arrow is drawn from the first to the second variant’s pie chart. Rings on the radial plot correspond to time in weeks; variants with later TOOs are placed more outward. Variants are arranged in different sectors (I-V), each of which is determined by the overall regional identity of the cells carrying the variants. Sectors are arranged from top (variants have more anterior destinations in the cortex) to bottom (more posterior destinations). For visual clarity, the insets at right show some of the early-rising variants. **(B)** Associations between estimated TOO and regional restriction as quantified by the regional restriction statistic (RRS) for each variant found in 2 or more cells in its corresponding lineage tree. Panels from top to bottom: The number of cells carrying each variant, the TOO estimates (in weeks) with 95% credible intervals, and the RRS computed for each variant (see Methods). The RRS range for germline variants is plotted as the gray band encompassing RRS=1 as a reference. Confidence intervals were constructed from bootstrapped estimates of RRS taken from sampling cells in clades of the lineage tree. **(C)** A schematic of the two main patterns that describe how variants (each belonging to a different sector) are dispersed across the cortex.

We grouped variants into five clusters distinguished by how cells carrying variants are spread out across the three regions. First, 18 variants across the two brains end up restricted to BA9 by PMW2 (**Fig. 2A, Cluster I**), suggesting that a subset of clones is isolated to the prefrontal cortex early in neurogenesis. Second and third, 22 observed variants end up primarily within the occipital cortex, either mostly restricted to BA17 (**Fig. 2A,Cluster V**) or split across BA17 and BA18 (**Fig. 2A, Cluster IV**) by PMW4-6. Nine of these 22 occipital variants were also detected at low cell fractions in BA9 at or before PMW6, suggesting that the exclusion of these variants from the frontal cortex is not complete before then. These three patterns represent the allocation of clones between the occipital and pre-frontal cortexes at different times. The fourth and fifth patterns represent the exclusion of clones away from BA17. We observed 41 variants excluded from BA17 over PMW1-3 that end up restricted to either BA18 by PMW6 (**Fig. 2A, Cluster III**) or to BA9 by PMW9 (**Fig. 2A, Cluster II**) separately from and more gradually than the early-BA9 variants (**Fig. 2A, Cluster I**). Given that cells in both clusters restrict to BA9, it is plausible that Cluster II may represent an intermediate state of Cluster I and that sampling more cells carrying sSNVs grouped into Cluster I may reveal more BA18/9 intermediate subclones.

For each variant, we defined its “regional restriction statistic” (RRS) (**Methods**) to quantify whether cells sharing it are spread out around a subset of regions (<1) or restricted to one region (>1), with germline or early-mosaic variants receiving ≈1 (equal representation across all regions). Over time and at increasingly lower cell fractions, sSNVs restrict to one of the three regions (RRS>1) (**Fig. 2B**). Despite the trend, two sSNVs at low cell fractions and arising between PMW5-10 remained distributed across two of the regions and depleted in one region (RRS<1). Although these variants resemble the pattern of highly-dispersed interneuron clones derived from the basal forebrain that have been described in other species (Cadwell et al., 2019; Subramanian et al., 2019; Tsai et al., 2012), their very low cell fractions make it difficult to capture enough cells carrying these variants to define the cell type associated with them.

In summary (**Fig. 2C**), using lineage trees and coalescent models, we infer that early cortical progenitors follow one of two paths. First, they may restrict to BA9 between PMW2-9 (GW12-19), with a possible transition through BA18 that leaves behind BA18-restricted descendants by PMW6 (GW16) (I/II and III). Second, progenitors may restrict themselves to the visual cortex and distribute across the BA17/BA18 border (IV and V). The exclusion of these clones from the frontal cortex is not complete until PMW6, and only after PMW10 (GW20) do these clones remain split between BA17 and BA18 (IV). BA17-restricted cells are descended from these posterior-restricting clones. From these paths, we infer that the distinction between frontal and posterior progenitors is made by PMW6. While a clone’s regional restriction generally increases with time and decreasing cell fraction, complete spatial restriction may not fully occur until late in the lineage (PMW10). We also predict that progenitors restricting towards BA17 will yield descendant cells with a more posterior orientation to cells descended from BA18-restricting progenitors, some of which will also show recent ancestry with cells in BA9.

### Low mosaic sSNVs disperse widely in the frontal cortex

Our results so far suggest a complex picture of how cortical clones can follow different routes towards populating the visual and prefrontal cortexes. To gain a more comprehensive picture of how cortical clones disperse over the course of development, we profiled a subset of sSNVs in a broader set of cortical areas and in non-cortical tissues. We conducted MIPP (Multiple Independent Primer PCR)-seq (Doan et al., 2021), which is capable of detecting a variant down to 0.1% MF (at >99% sensitivity), well below the 2% MF threshold of 210X WGS. We analyzed 59 sSNVs (discovered from WGS and where probes could be successfully designed) across 25 cortical regions and 5 non-cortical regions (**Fig. 3A, Fig. S3A**, and **Table 1**). We previously identified clonal clades in UMB4638’s frontal cortex using four of these sSNVs (labeled as “A-early,” “C-early,” “B-early,” and “B-later”) (Huang et al., 2020). With MIPP-seq we could track these and other rarer sSNVs across a wider area, allowing us to delve deeper into the clonal structure of the frontal cortex (**Fig. 3B**). We refer to MFs of ⪅1% as “ultra-low” mosaic and 1-3% as “low” mosaic (**Fig. 3**,**Fig. 4, Fig. S4** and **Table 3.1**).

**Figure 3:**
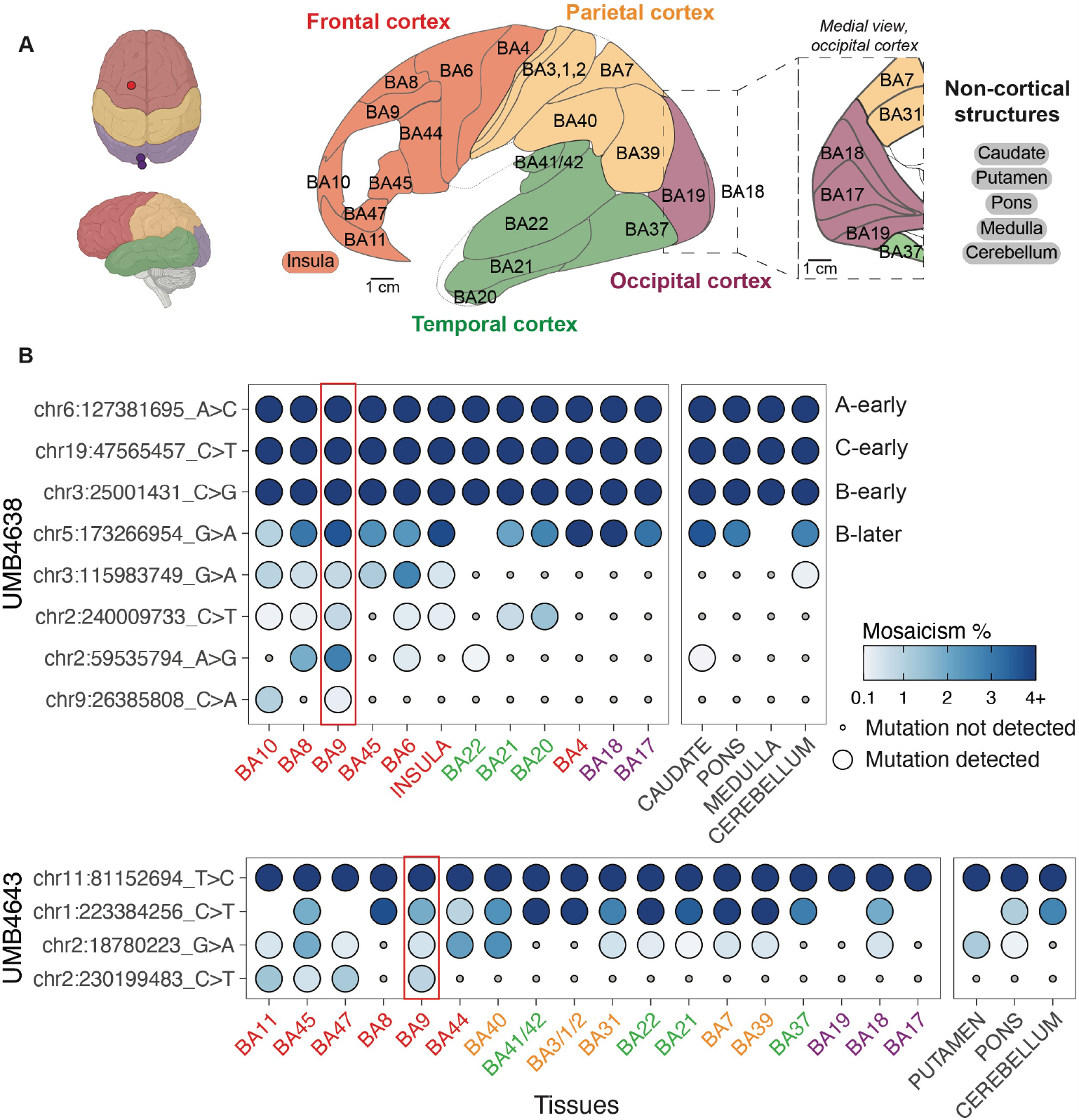
Clonal sSNVs identified in frontal cortex (BA9) distribute across cortical regions, with modest regional restrictions, at low mosaic fraction. **(A)** The brain regions that were sampled for studies of the spread of sSNVs using MIPP-seq. Cortical lobes from which Brodmann areas/cortical structures were taken are colored (frontal, red; parietal, yellow; temporal, green; and occipital, purple; non-cortical, black). Regions separated by slashes (BA3/1/2 and BA41/42) were studied together. “A-early,” “C-early,” “B-early,” and “B-later” are four variants that we previously had studied in the frontal cortex for UMB463818. **(B)** The cortical distribution of a sSNV originally detected in BA9 (red rectangle) from UMB4638 (top) and UMB4643 (bottom). Tissues are arranged (left to right) in anterior to posterior cortical section ordering, and are color highlighted based on the scheme in **(A)**. Non-cortical tissues are listed on the right. Mutations are ranked from broadest to least present across the tissues, followed by average mosaicism across samples.

**Figure 4:**
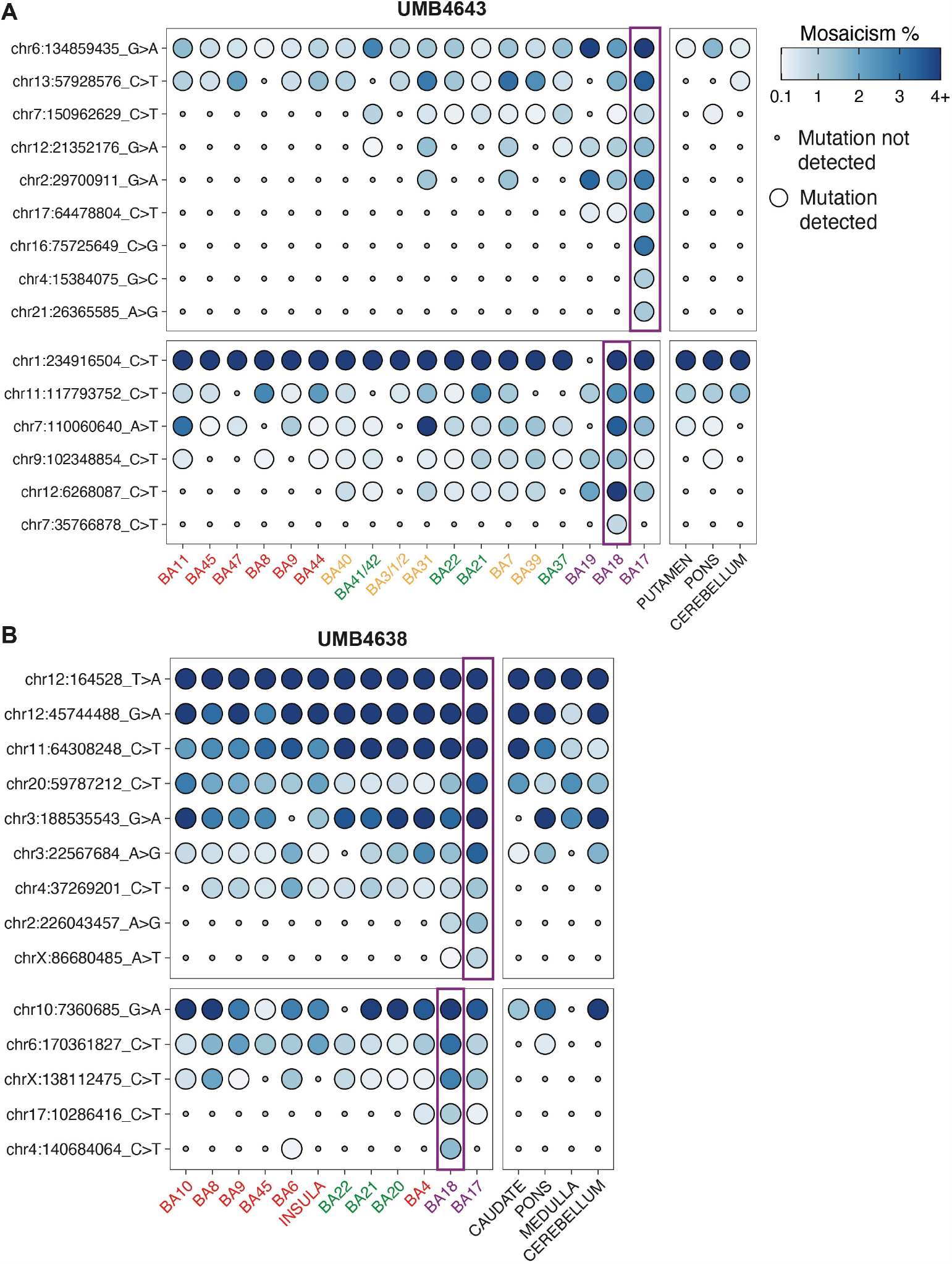
The spatial distributions of the rarest sSNVs initially discovered within the occipital cortex. The presence of select sSNVs initially discovered in BA17 or BA18 within the original discovery site (purple rectangles) and across adjacent and distant cortical regions. sSNVs show either regional restriction **(A)** or distinct patterns of lobe restriction after crossing the BA17/18 border **(B)**. Tissue and mutations from UMB4638 and UMB4643 were assessed and are colored as in Fig. 3.

MIPP-seq analysis showed a subset of clones with wide regional dispersion even at low MFs. Low and ultra-low mosaic sSNVs discovered within BA9 showed wide distribution across the frontal cortex and even in the temporal cortex of the same hemisphere but were typically not detected in the occipital cortex. For example, in UMB4638, chr9:26385808 was detected in two adjacent cortical areas: BA9 (0.22% MF at ODS) and BA10 (1.02% MF). Similarly, in UMB4643, chr2:230199483 is detected all throughout the frontal cortex beyond BA9. Two other variants in UMB4638 (chr2:240009733, chr2:59535794), originally discovered at low MFs in BA9 (0.77% and 2.85%, respectively), were detected as far away as the temporal lobe (the insula and BA20/21/22/37/41/42) at MF ≤ 1.41% (**Fig. 3B**). In our WGS analysis, we found evidence for a significant number of region-specific sSNVs in BA9 present in >1% of total cells (**Fig. S1C**). By following some of these variants with MIPP-seq, we demonstrate that even late-rising variants initially discovered in BA9 may be dispersed into neighboring and distant territories. The frontal cortical lineages marked by these variants may be significantly intertwined with those from other regions.

### Low mosaic sSNVs in visual cortex tend to distribute asymmetrically at the BA17/*BA18 border*

In contrast to their wide dispersion across frontal cortex, low and ultra-low mosaic sSNVs (≤ 3.1% MF) are asymmetrically spread across the BA17/18 border, showing stronger regional or posterior-lobe restriction in BA17 compared to the adjacent BA18. For example, three sSNVs discovered in BA17 of UMB4643 with MFs of 1.1-3.1% (chr16:75725649, chr4:15384075, chr21:26365585) were detected only in BA17, whereas a single sSNV (chr7:35766878; 0.77% MF) was detected only in BA18 (**Fig. 4A**). Although the sample size is small, these observations are consistent with the results observed in WGS in which BA17 variants showed more region-restriction than BA18.

On the other hand, some low mosaic sSNVs (≤2.14% MF) were not limited to their original discovery sites in WGS at BA17 or BA18 but crossed over into the neighboring regions. In UMB4638, chr17:64478804 was originally found in BA17 (≤ 2.14% MF) but also seen in BA18 (0.21% MF) and BA19 (0.30% MF), all in the occipital lobe (**Fig. 4A**). Two variants in UMB4638, chrX:86680485 and chr2:226043457, were discovered in BA17 (present at respective MFs of 0.86% and 1.54%) but also detected in BA18 (respective MFs of 0.10% and 0.84%) (**Fig. 4B**). These observations are consistent with the inferences from our lineage trees suggesting significant intermingling of occipital lobe lineages even late into cortical development.

The MIPP-seq data also captured a prediction initially made in our lineage analysis: the spread of BA18-discovered sS-NVs in more anterior regions of the cortex compared to those containing BA17-discovered sSNVs. Whereas sSNVs originally discovered in BA17 or BA18 (MFs 1.61–7.98%) were detected in the occipital, parietal, and/or temporal cortices at maximum MFs of 3.34% (**Fig. 4B**), the domain of BA18-discovered variants appeared more anterior to the domain of BA17-discovered variants. sSNVs originally discovered in BA18 (e.g., chr17:10286416 and chr4:140684064) at low MFs (1.11%-1.60%) were detected at lower MFs (0.15-0.41%) in posterior frontal lobe (BA4 and BA6, **Fig. 4B**). On the other hand, one sSNV originally discovered in BA17 (chr12:21352176) at a low MF (1.61%) was found beyond the occipital cortex down to 0.11% MF across the temporal and parietal cortex but at regions more posterior to the boundary of BA18-originating variants (BA37, BA41/42, BA7, BA31). Additionally, this specific sSNV was not detected in the frontal lobe, unlike similarly rare sSNVs from BA18. This observation replicates the inference made in our lineage analysis (**Fig. 2A**) that BA18-discovered clones may have more anterior spread across the cortex (as in Cluster II) than BA17-discovered clones (as in Clusters IV/V).

In contrast to the overarching trend of low-mosaic sSNVs showing modest regional restriction, a subset of low-mosaic sSNVs (at≤ 2.81% MF) showed no evidence for regional restriction at all, appearing all across the hemisphere but not in subcortical and non-cortical brain tissues, a pattern reminiscent of inhibitory interneuron clones described in animal models (Cadwell et al., 2019; Subramanian et al., 2019; Tsai et al., 2012). In UMB4643, some sSNVs showed an irregular, spotty pattern within the cortex, non-cortical brain tissue, and the putamen (chr9:102348854 and chr13:57928576; average 0.42–1.14% MF across tissues, **Fig. 4**). These variants represent exceptions to the gradual regional restriction of most variants, although determining their corresponding cell types (e.g., whether they represent inhibitory neuron clones) would require deep targeted capture of these extremely low frequency clones.

In summary, tracking a handful of variants with MIPP-seq replicates three trends seen in our WGS and lineage tree analysis: sSNVs discovered in BA17 tend to remain regionally restricted, sSNVs discovered in BA18 tend to intermingle with regions as far as in the frontal cortex, and sSNVs discovered in BA9 are more likely to be restricted to the anterior end of the brain while dispersing widely within the frontal lobe. Although MIPP-seq does not establish overarching trends as do our WGS and lineage analysis; it highlights nuances in the clonal structure of the cortex through a deep dive of a subset of variants. MIPP-seq analysis over the entire cortex further suggests that for the rarest low mosaic clones from BA9, BA18, and BA17; lower-MF sSNVs show more restricted presence in cortical areas or specific lobes in the cerebral cortex (Breuss et al., 2022; Cadwell et al., 2020). However, the detection of low and ultralow-MF clonal sSNVs in adjacent and distant regions suggests that human cortical lineages do not remain laterally constrained throughout most of development.

### Low mosaic sSNVs in visual cortex tend to distribute asymmetrically at the BA17/BA18 border

Although coverage of sSNVs in single-nucleus (sn) RNA-seq and snATAC-seq is generally sparse (Bizzotto et al., 2021; Petti et al., 2019), the presence of pre-specified variants can be verified in a fraction of cells. We collected snRNA-seq and snATAC-seq data from 10X Chromium libraries of DAPI-sorted or NeuN+-enriched FACS-sorted cells from BA17, BA18, and BA9 across UMB4638 and UMB4643 and determined cell types using markers determined by the Allen Brain Atlas39 (**Methods**). We retained 72,839 nuclei across 15 snRNA-seq experiments and 9,125 nuclei across 2 snATAC-seq experiments (Fig. 5A,B). We sought to profile 350 candidate sSNVs in UMB4638 and 306 in UMB4643, including those used for lineage tracing as well as additional variants subjected to amplicon validation (**Methods**; **Table S1.1-1.5**). Each cell had on average 1-4 reads of coverage per site (**Fig. S5A-B**), and the overall cell fraction of a given variant correlated with its average WGS AAF across regions (**Fig. S5C**). Although only 15% of our assessed variants had sufficient read coverage to produce at least one read of a mutant allele (**Fig. S5D**), we found variants in each major cell type in our dataset (**Fig. S5E**) and identified numerous subsets of cells that shared sSNVs (**Fig. S5F**).

**Figure 5:**
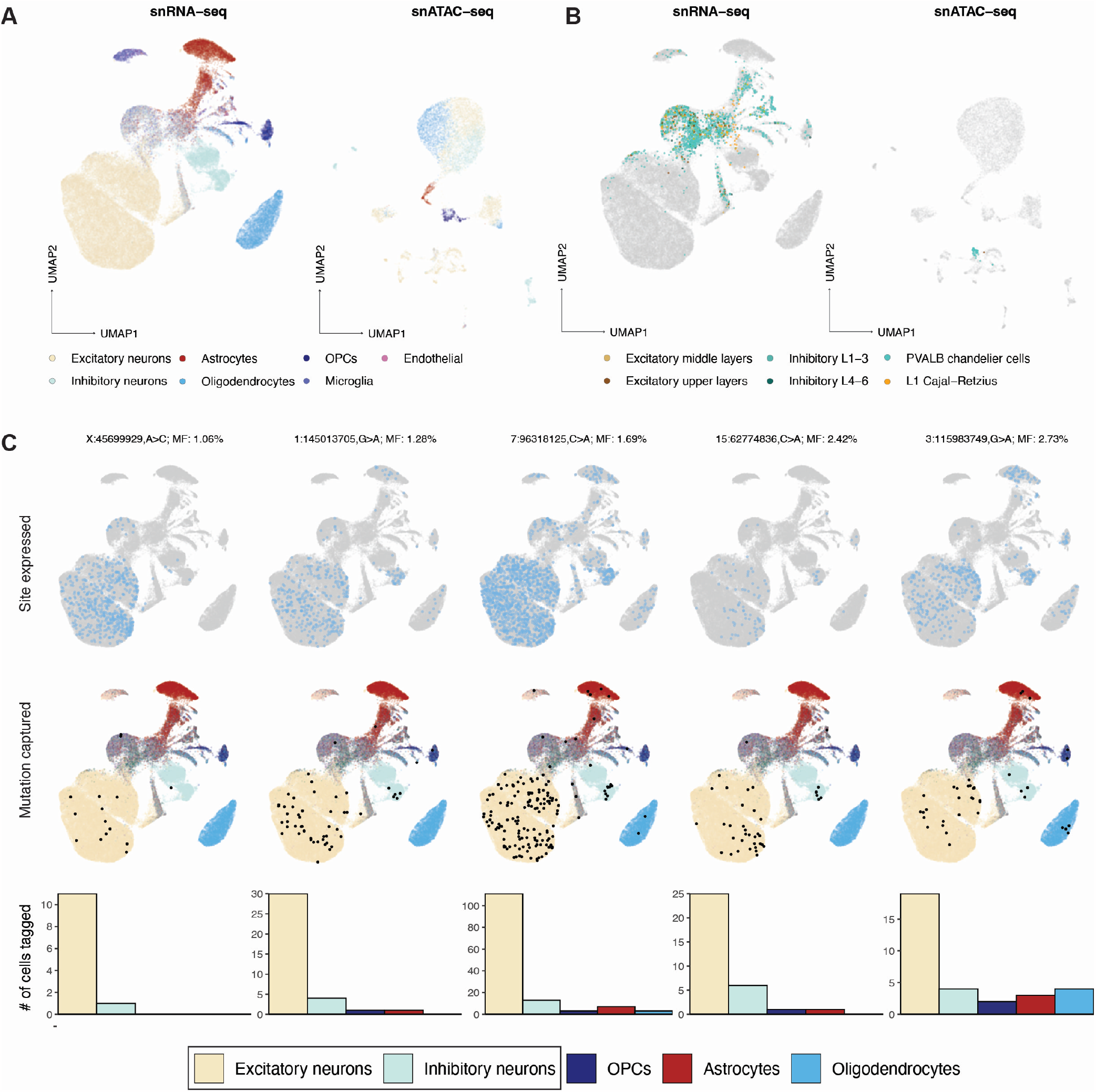
Low-mosaic sSNVs are shared by both excitatory and inhibitory neurons. **(A and B)** Low-dimensional UMAP projections of 72,839 nuclei from snRNA-seq and 9,125 nuclei from snATAC-seq, both collected from 10X Chromium assays. Major cell types **(A)** and more specific neuronal subtypes **(B)** are colored. **(C)** Five select sSNVs at low mosaicism (¡3%) and the cell types in which they are detected. Top panels: cells are marked in light blue if they express the corresponding sSNV’s locus. Middle panels: cells are marked in black if they express the mutant allele; otherwise, they are colored by their assigned cell type. Bottom panels: bar plots marking the number of cells in each type that express the mutant allele.

Strikingly, a number of these low-mosaic sSNVs are found in both excitatory and inhibitory neurons (EN and IN; **Fig. 5C**), two distinct neuronal subtypes that have been reported to arise from anatomically distinct progenitors in rodents. Existing studies in mice suggest that INs migrate into the cortex after arising from the medial or caudal ganglionic eminences (MGE and CGE) deep within the developing brain, separately from the dorsal cortical progenitors of the ENs closer to the outside surface (Lim et al., 2018; Bandler et al., 2017). However, several reports suggested a dorsal source of INs in the human neocortex (Al-Jaberi et al., 2015; Clowry, 2015; Cunningham et al., 2013; Fertuzinhos et al., 2009; Jakovcevski et al., 2011; Letinic and Rakic, 2001; Letinic et al., 2002; Petanjek et al., 2009; Rakic and Zecevic, 2003; Radonjić et al., 2014), and a recent report using viral lineage tracing of human progenitor cells in rodent xenografts (Bandler et al., 2021) suggested that cortical ventricular zone progenitors produce proportions of 67-85% excitatory to 4-11% INs (Delgado et al., 2021). If some INs and ENs share a direct dorsal progenitor, then these neurons would likely share sSNVs present at lower cell fractions, given that dorsal progenitors are only a subset of the cells in the developing brain.

To systematically analyze the relatedness of ENs and INs to each other as well as to other brain cell types, we measured the AAFs of sSNVs shared across cell types. To differentiate the early- and late-diverging cell types, we reasoned that the AAF of variants shared in two cells will be higher if the cells diverged earlier in developmental time (if multiple variants are shared, we take the minimum AAF) (**Fig. 6A**). Using low AAF (<5%) variants that are likely to be informative for brain-specific hierarchies, we inferred the relatedness of neurons to basic cell types in the brain (**Fig. S6A**). We observe only one variant shared between microglia and excitatory neurons at a low AAF ( 5% AAF) but dozens more amongst neurons and macroglia (astrocytes, oligodendrocytes, and OPCs) at lower AAFs (1-2%) (**Fig. 6B**). We were surprised to find sSNVs at low AAFs (<1%) shared not only within ENs (E/E) or within INs (I/I) but also between ENs and INs (E/I). We confirmed that both non-mutant and mutant UMIs were expressed for each variant in both neuronal subtypes, confirming that there is no confounding due to transcript dropout and that we have sufficient coverage and sensitivity (**Fig. S6B**). The minimum AAF of E/I sSNVs was lower than that of sSNVs shared between neurons and different macroglial subtypes. As macroglia may arise from the same radial glial progenitors as neurons (Rakic, 2009; Noctor et al., 2001), the lower AAFs of E/I sSNVs compared to those shared between neurons and macroglia suggest a common ancestor of E/I neurons present later than that of neurons and macroglia. Expanding our view to all sSNVs shared amongst neurons, we found that E/E, I/I, and E/I sSNVs showed surprisingly similar AAF distributions (**Fig. 6C**). Overall, our analysis indicates early divergence of neurons from microglia, late divergence from macroglia (astrocytes, oligodendrocytes, or OPCs), and a later divergence amongst neuronal subtypes, including E/I clones.

**Figure 6:**
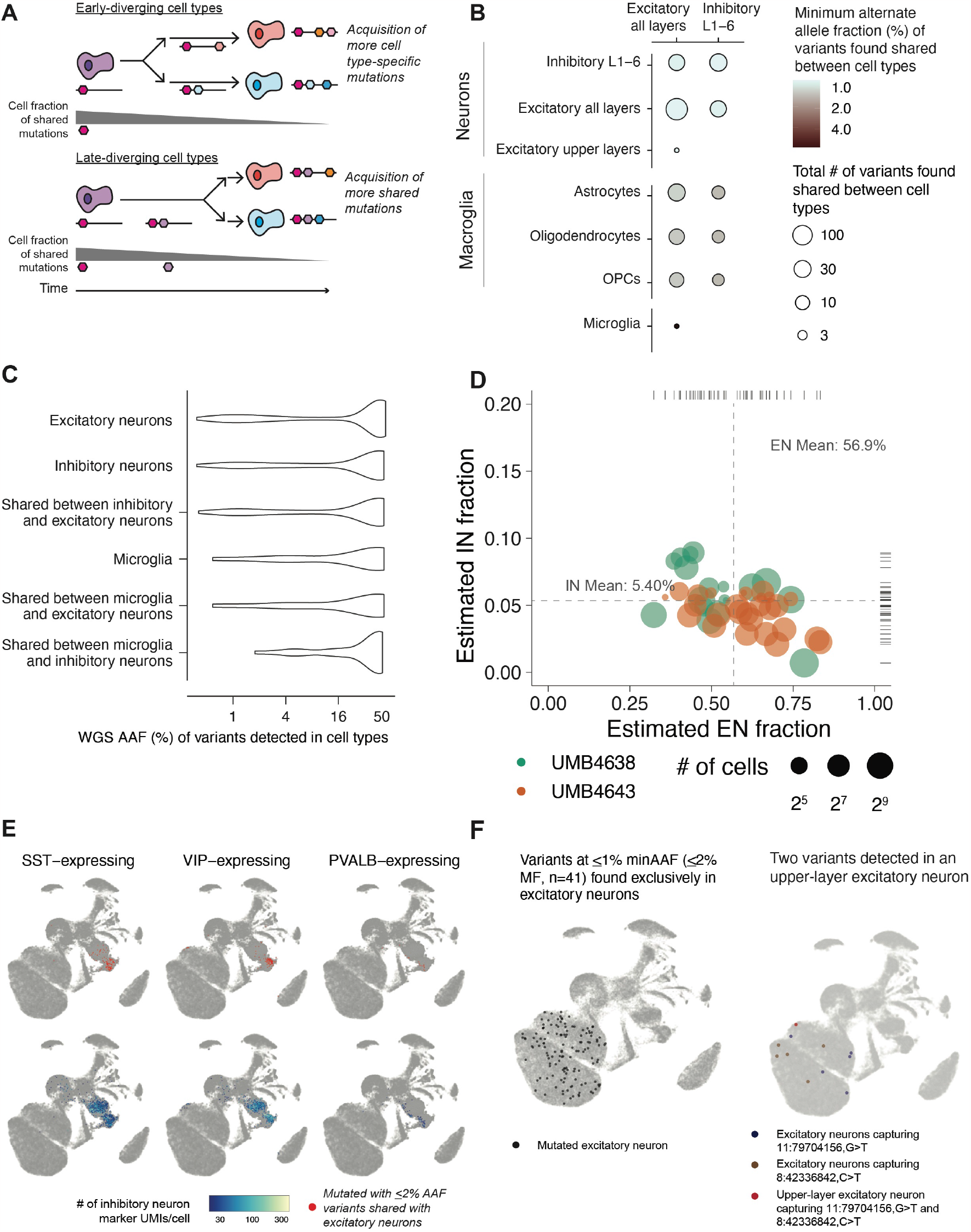
Properties of low-mosaic excitatory and inhibitory clones in the dorsal cortex. **(A)** A schematic depiction of the motivation behind the minimum-AAF statistic, used to assess if a variant shared between two cell types occurred in a progenitor before an early or late divergence of the cell types. **(B)** Number and minimum AAF (from region-averaged WGS) of variants found detected across cell types annotated within the 10X data. Estimates were aggregated across all variants detected with 10X data from both brains. **(AC** WGS AAF distributions of variants found detected in excitatory neurons, inhibitory neurons, and microglia. **(D)** Empirical Bayes estimates of the proportion of excitatory and inhibitory neurons found in subsets (as depicted in (B)). Only subsets with ¿ 10 cells were analyzed. **(E)** UMAP annotations of inhibitory neurons (marked in red) sharing variants at ¡2% AAF (¡4% MF) with excitatory neurons (not marked), along with the expression level (number of UMIs) of different types of inhibitory neuron markers. **(F)** UMAPs with variants marked for variants found exclusively shared amongst excitatory neurons (all layers) (left panel) and for two variants shared between an upper-layer neuron and subsets of all-layer excitatory neurons (right panel).

If late E/I progenitors exist in vivo in the cortex, they may be the source of low-AAF E/I sSNVs and produce the two neuronal types at consistent ratios. To further investigate this hypothesis, we estimated the ratios of excitatory to inhibitory neurons sharing low-MF E/I sSNVs. Using Empirical Bayes estimation (**Methods**), we found this ratio (E/I) to be approximately 10:1 (EN:IN of 56.9%:5.40%; 95% CIs of 39.0-69.1% and 4.14-7.6%, respectively; Figure S6C,6D), surprisingly consistent with ratios of these cells seen in xenograft experiments (Delgado et al., 2021). Inhibitory neurons sharing variants with excitatory neurons showed diverse gene expression profiles, predominantly those similar to VIP-expressing, PVALB-expressing, or SST-expressing inhibitory neurons as determined in the Allen Brain Atlas (Hodge et al., 2019) (**Fig. 6D, Fig. S6D-E**). From 32 E/I sSNVs at <4% AAF (¡8% MF), we found that some E/I clones are regionally restricted (as some ENs have been thought to be), while others are broadly dispersed across the cortex (as some INs have been thought to be) (**Fig. S6D**). We could not definitely establish if inhibitory neurons expressing MGE or CGE markers differed in the MFs of E/I sSNVs (**Fig. S6F**).

### Inferring the timing of specialized neuron types

In addition to studies of E/I clones, analyzing the AAFs of sSNVs shared amongst neuronal subtypes allowed for the closer analysis of EN-exclusive clones. We identified 41 E/E variants at <1% AAF in multiple cortical layers and detected in no other cell types (**Fig. 6F**). We also found two variants at <1.5% AAF shared between one excitatory upper-layer neuron and other excitatory neurons but no other cell types (**Fig. 6F**). Given their presence in the same upper-layer neuron, these two variants likely occurred within the same population of excitatory neurons, but the low per-cell coverage from Chromium assays means that not all excitatory neurons have enough coverage to report the variant.

For one of the upper-layer EN sSNVs and two of the E/I sSNVs, we could estimate TOOs due to overlaps with our lineage tree (**Fig. S6H**). The upper-layer EN sSNV (chr8:42336821) had a TOO of PMW3.5 (≈GW13.5, if the MRCAs of the single cell lineage trees are assumed to have arisen near the onset of neurogenesis at≈ GW10). Despite its relatively early TOO, this variant remains largely restricted to BA9 at a low MF with a much scarcer presence in BA18 and undetectable in BA17. This particular variant may support the presence of upper-layer restricted progenitors in the human cortex yielding spatially restricted neurons, an observation first seen in mouse models (Franco et al., 2012; Gil-Sanz et al., 2015; Eckler et al., 2015). The first E/I sSNV (chr3:115983749) had an estimated TOO of PMW9.5 (≈GW19.5), was mapped to a BA18/BA9-distributed population in the single-cell lineage tree, and shows a similar regional distribution in WGS data. The second E/I sSNV (chr3:65583407) had an estimated TOO of PMW10.7 (≈ GW20.7) and was also mapped to BA9 and BA18 in the lineage tree, but it was present at higher MFs across all three regions. This discrepancy may arise from the more unbiased sampling of tissue in bulk WGS and the greater potential for single-cell dropout from single-cell sampling. These timing estimates of E/I sSNVs may hint at a late generation time for ENs and INs from common dorsal progenitors.

### Targeted DNA/RNA analysis further defines cell type relationships

To confirm patterns of divergence of excitatory and inhibitory lineages, we used Parallel RNA and DNA analysis after Deep sequencing (PRDD-seq) as a complementary method of targeted analysis of multiple RNA markers of cell type and multiple sSNVs within the same single cells (Huang et al., 2020) (**Fig. S7A**). We first applied PRDD-seq on variants previously identified from BA9 (“Clade A” (Huang et al., 2020)) and integrated the data with topographic mapping from MIPP-seq (**Fig. 3B**). Early variants in Clade A (A1-4) mapped to both excitatory and inhibitory cell clusters, including the four evaluated inhibitory neuron subtypes (**Fig. 7A-B** and **Fig. S7**). Lower-mosaic sSNVs in Clade A (A5) are increasingly restricted towards excitatory neurons in upper layers (**Fig. 7A** and **Table S6.1**), consistent with our findings of upper-layer excitatory neurons sharing low-MF sSNVs with only other excitatory neurons (**Fig. 5D-E**). Within Clade A, sSNV A5b (chr17:48665916) represents a later-occurring mutation (**Table S6.1**) that is limited to later-born excitatory neurons in middle and upper cortical layers (**Fig. 7B**) but nonetheless disperses across >5.2 cm in the frontal cortex across (in rostro-caudal direction) BA10, BA9, BA8, and BA6 (**Fig. 7C**). Restriction of A5b to middle- and upper-layer excitatory neurons matches the inside-out sequence of excitatory neuronal cell types in the cerebral cortex seen in several animal models, in which deep layer neurons are formed first, and the most superficial excitatory neurons are formed last (Harwell et al., 2015; Mayer et al., 2015). Overall, these observations confirm our previous inferences that late-occurring variants arising after generation of layer 6 neurons can nonetheless spread widely across many centimeters and Brodmann areas.

**Figure 7:**
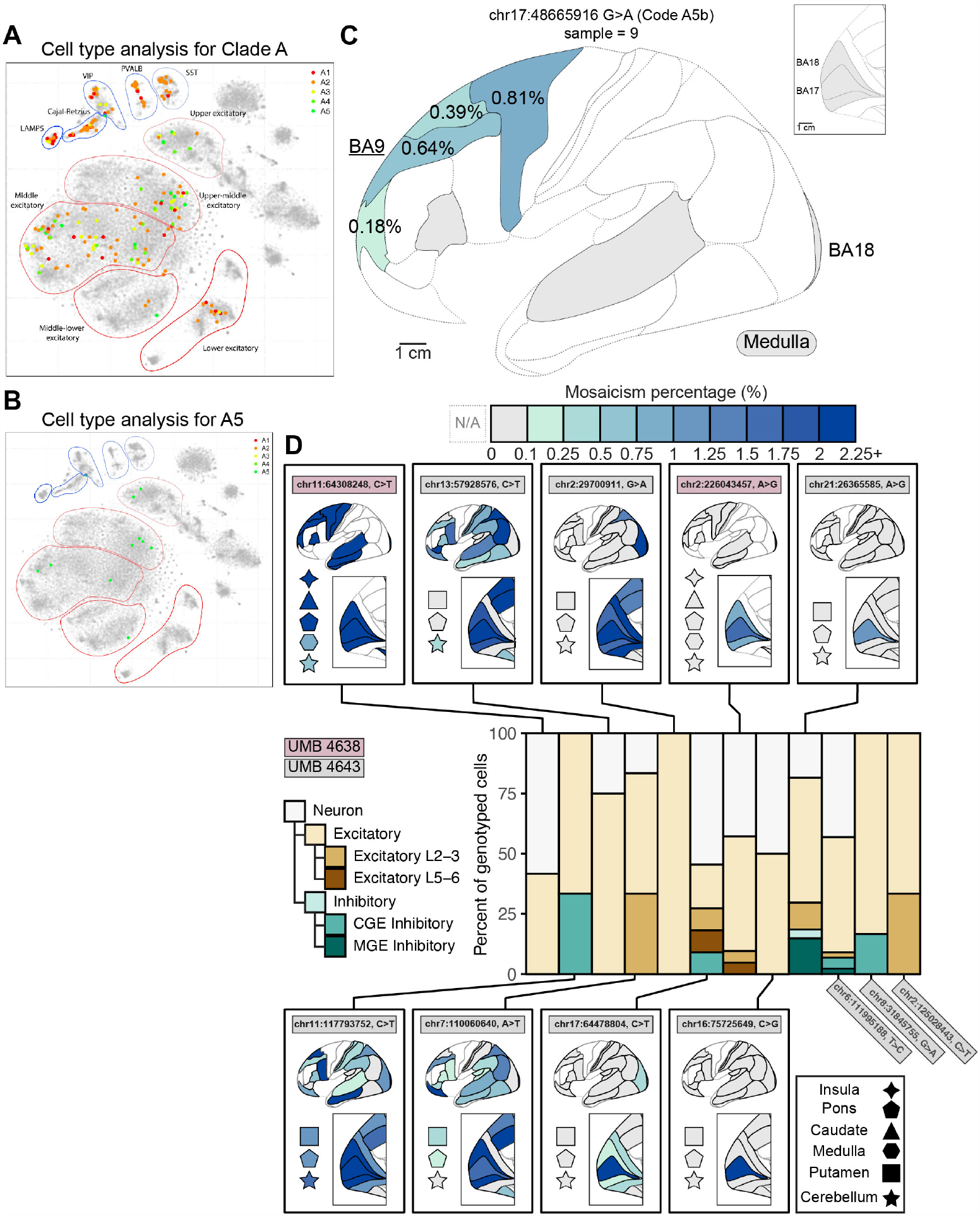
PRDD-seq further reveals the clonal topography and relatedness of excitatory and inhibitory neuron subtypes. **(A-B)** PRDD-seq-evaluated sSNVs across three clades in UMB4638 distribute throughout different excitatory and inhibitory subtypes, with restriction to middle- and upper-layer excitatory neurons seen in later-occurring sSNV. **(A)** Low-dimensional representation of the transcriptomic profiles of single neurons carrying sSNVs that were previously characterized in lineage studies of BA9 (Clade A18). Neurons were collected and analyzed with PRDD-seq were found to carry sSNVs from successive generations within Clade A (earliest, A1; latest, A5) and their marker profiles are used to project them (marked as solid points) onto a t-SNE representation of snRNA-seq data from UMB4638 and UMB4643 (Methods). Clusters of excitatory and inhibitory neuron subsets are indicated. **(B)** Cell type breakdown for sSNVs in A5. sSNV A5 is found in middle- and upper-layer excitatory neurons but is mostly absent from deep layer excitatory neurons and from inhibitory neurons. **(C)** Example of a late-occurring A5 sSNV (A5b) that was identified in BA9 (underlined), is limited to middle-upper layer excitatory neurons, and marks a neuronal clone distributed widely ( ≥ 5.2 cm) and at ¡1% MF across frontal cortical areas. The topographic distance between the furthest sampled and detected cortical areas is a minimum total distance of 5.2 cm, but likely 6.2 cm since one cortical section was unavailable for sampling and measurement. **(D)** Regional presence of ultra-low mosaic sSNVs and the neuronal subtypes harboring them as assessed by MIPP-seq. PRDD-seq reveals mosaic sSNVs in UMB4638 (brown) and UMB4643 (gray) limited to BA17 (chr21:26365585, A¿G, chr16:75725649, C¿G), BA17 and BA18 (chr2:226043456, A¿G), occipital lobe (chr17:64478804, C¿T) or occipital and neighboring parietal lobe areas (chr2:29700911, G¿A). The sSNVs are mainly found in excitatory neurons, though some are found in inhibitory neurons as well consistent with a dorsal source of some inhibitory neurons 20. PRDD-seq analysis was conducted completed in single neurons isolated from BA17 from each individual.

We then applied PRDD-seq to 14 sSNVs of UMB4638 and UMB4643 for which we could successfully design PRDD-seq primers. We could confidently isolate, genotype, and classify the cell types for neurons carrying 12 of these sSNVs, 9 of which also had cortex-wide topological information from MIPP-seq. Analysis of BA17 from UMB4638 and UMB4643 showed several topographically restricted sSNV recovered in excitatory neurons but also detectable in inhibitory populations (**Fig. 7D** and **Table S6.2**). A sSNV (chr21:26365585) restricted to a single cortical area (BA17) was positively genotyped in 17 excitatory and 5 inhibitory neurons, with 5 other identified neurons not further assignable to subtype (**Fig. 7D**). Similarly, sSNV chr17:64478804 ( ≤ 2.14% MF) was detected in all occipital lobe areas (BA17, BA18, and BA19) and in 11 neurons (4 excitatory, 1 inhibitory, and 6 neurons not further classified; Figure 7D). Other sSNVs detectable in excitatory neurons (chr16:75725649, chr8:31845755, and chr2:226043457) genotyped in various excitatory neuron subsets were found in other unassignable neurons. Low mosaic SNVs showing widespread or non-regional or lobe-restricted patterns (chr11:64308248; chr11:117793752; chr13:57928576; chr7:110060640; and chr2:29700911) were also commonly seen in excitatory neurons (**Fig. 7D**), but as can be seen with chr11:117793752 (**Fig. 7D** and **Table S6.2**) may contain inhibitory neurons, confirming that sSNVs marking clones dispersing across several cortical boundaries at low MF represent inhibitory neurons, but the pattern reinforces our observation that some excitatory neuron clones may also disperse across cortical boundaries. Since inhibitory neurons are both less common and expected to be extremely widely dispersed, our single-cell methods are relatively insensitive to sampling clonally related inhibitory neurons. Nonetheless the consistent co-occurrence of E/I sSNV with two methods supports the presence of a shared E/I progenitor relatively late in neurogenesis in the dorsal cortex, consistent with previous xenograft lineage tracing (Wichterle et al., 1999).

## Discussion

By using somatic mutations as markers of cell lineage in the human cerebral cortex, we find major aspects of cell lineage that are quite unlike anything described in animal models to this point. Three methods—deep WGS, coalescent analysis of single cell lineages, and MIPPseq analysis of cortical dispersion—all suggest that low mosaic sSNVs (MFs <1%) disperse widely across cortex in general, but show prominent non-uniformities across the BA17/18 border. BA17 harbors more regionally-restricted sSNVs than BA18 (**Fig. 1C**), likely reflecting regional differences in proliferation, and clonal intermingling appears somewhat restricted across this border. In addition, two methods combining DNA analysis with single-cell transcriptomics suggest that widely dispersed clones contain many excitatory neurons and reveal clones of E/I neurons likely arising from the dorsal proliferative region in vivo (Cai et al., 2013; Radonji ćet al., 2014; Letinic et al., 2002).

### Clonal patterns at the BA17/18 border

Our data mirror other findings in primates that the BA17/18 border shows somewhat limited clonal dispersion. Studies of non-human primates show a sharp change in patterns of proliferation in the subcortical proliferative zones beneath this border, as well as relatively constrained patterns of radial migration of BA17 neurons (Dehay et al., 1993; Smart et al., 2002; Lukaszewicz et al., 2005; Cortay et al., 2020). In single-cell tracing studies of the E78 subplate in macaques that underlies BA17/18, BA17 showed more radial trajectories of migrating supragranular neurons than BA18 (Cortay et al., 2020), in addition to a unique dependence on visual inputs for its proper development. The frequently identified high neuronal density observed in BA17 compared to BA18 and other cortical areas (Greig et al., 2013; Collins et al., 2010) may reflect the observed higher tendency for sSNVs identified in BA17 to be restricted to BA17, compared to BA18 or BA9 where sSNVs are not as commonly restricted to the region where they were discovered. To reconcile our data with these findings, we propose that in addition to dispersion patterns seen elsewhere in the cortex, clones at the BA17/18 border also show BA17-restricted increase in proliferation leading to clonal expansion, potentially regulated by signals from thalamic fibers, and likely involving predominantly upper-layer neurons. A larger amount of local proliferation of neuronal precursors would give rise to a higher neuronal density and increased clonality marked by sSNVs detectable at our MF threshold. Alternatively, our data are also consistent with a model in which the border unevenly allocates progenitors between BA17 and BA18 and restricts BA17-derived clones from crossing over into the rest of the cortex, while BA18-derived clones are not as restricted and thus end up as far as the frontal cortex.

### Widespread clonal distributions of excitatory neuron clones in cerebral cortex

Our findings confirm earlier reports of broad dispersion and intermingling of clonal progeny in the human frontal and lateral cortex (Lodato et al., 2015; Breuss et al., 2022; Bizzotto et al., 2021; Huang et al., 2020; Coorens et al., 2020; Evrony et al., 2015; Lavdas et al., 1999; Fasching et al., 2021), but they show for the first time that these widely dispersed clones include excitatory neurons. While previous studies have shown that specific excitatory neuron clones intermingle within a single cortical column (Huang et al., 2020), we observe that excitatory neuron clones present in all cortical layers typically encompass most or all of the cortical surface. Excitatory neuron clones disperse across multiple cortical areas at MFs as low as <1%, especially in frontal lobe, although previous limited analyses have suggested that later clonal events can show more limited dispersion across cortex at MFs <<1% (Evrony et al., 2015). Although the latest-generated sSNVs are hard to recover, we have also found that sSNVs at <1% MF with topographic restriction are often limited to neurons in middle-to-upper cortical layers, although confirming this finding will require analyzing more samples. Nonetheless, this extreme level of clonal intermingling for even rare, late-rising excitatory clones has major consequences for models of clonal structure in humans (Li et al., 2012).

The wider dispersion of excitatory neuron-generating clones in human cortex contrasts with the more coherent clonal patterns reported in the rodent cortex (Gao et al., 2014; Ware et al., 1999; Llorca et al., 2019), though limited reports in larger-brained mammals like ferret and nonhuman primate hint at wider clonal dispersion in these species as well (Ware et al., 1999; Reillo et al., 2017; Gertz and Kriegstein, 2015; Reid et al., 1997). It is unclear for now whether the human reflects a scaled-up version of similar mechanisms in non-primates or shows newly evolved mechanisms. One implication of the widespread clonal dispersion in humans is that pathogenic somatic mutations affecting human neuronal clones, such as seen in focal cortical dysplasia, may be scattered widely across the cortex, potentially beyond the borders of visible cortical lesions caused by these mutations (Fauser et al., 2004, 2015; Hamiwka et al., 2005).

### In vivo evidence for common dorsal progenitors of cortical excitatory and inhibitory neurons

We identified sSNVs shared by excitatory and inhibitory interneurons (E/I variants) in spatially restricted patterns that are characteristic of dorsally derived excitatory clones. This observation suggests a dorsal source of cortical interneurons in vivo, as has been suggested before and shown recently for human cells grown as xenografts (Delgado et al., 2021). This pattern contrasts sharply with mouse, where interneurons appear to be exclusively derived from subcortical sites (Miyoshi et al., 2010; Wonders and Anderson, 2006). While interneuron formation certainly occurs at subcortical sites in humans (Hansen et al., 2013; Ma et al., 2013), dorsal sites seem to be a significant source of them as well. Past studies suggest that the global E/I ratio in humans is approximately 7:3, compared to 3:1 in marmosets (non-human primates) and 8.5:1.5 in mice (Loomba et al., 2022; Bakken et al., 2021; Džaja et al., 2014). The 10:1 ratio of E/I cells in mixed E/I cortical clones we observed suggests that a substantial minority of the human cortex INs are dorsally derived. The absence of similar information from other mammalian species leaves open whether this dorsal interneuron source is a primate-derived addition, or whether mice selectively lost this dorsal source. Additionally, while we interpret our data as suggesting a dorsal origin for some interneurons, it does not exclude a ventral progenitor migrating from the ganglionic eminences into the cortex before producing E/I progeny. Future work must efficiently discover new sSNVs in interneurons to trace the separate lineages of dorsal and ventral progenitors.

### Limitations

Lineage studies in humans remain limited by small sample sizes, reflecting the lack of high-throughput, droplet-type methods to allow simultaneous analysis of DNA and RNA in single cells. The vastness of the human cortex also remains an obvious challenge for systematic description. Our retrospective method also does not allow direct determination of where neurons are formed, only their final location. Furthermore, present methods to identify sSNV, such as deep WGS, or single cell WGS, have far higher sensitivity to identify sSNV that occur early in development, prior to neurogenesis, and much lower sensitivity to identify the “late branches” of neurogenesis which are in principle the most informative to examine regional and cell type decisions. This lack of sensitivity especially impacts the analysis of interneuron lineages, which in animal models are highly dispersed (Letinic and Rakic, 2001; Radonjićet al., 2014; Harwell et al., 2015; Mayer et al., 2015; Wichterle et al., 1999), and which likely correspond to the outlier sSNV that we observe with cortex-wide dispersion at very low cell fraction. The tremendous clonal dispersion of these rarer cell classes means that sSNV marking them will be present at exceedingly low cell fractions that will challenge the sensitivity of present methods. Systematic application of newer duplex sequencing methods (Abascal et al., 2021; Xing et al., 2021) promise to improve sensitivity for these late-occurring variants.

## Acknowledgements

Human tissue was obtained from the NIH NeuroBioBank at the University of Maryland, and we thank the donors and their families for their invaluable donations for the advancement of science. We thank R. Mattieu, K. Brownstein, J. Li, the Flow Cytometry Facility in Boston Children’s Hospital, Boston Children’s Hospital Intellectual and Developmental Disabilities Research Center Molecular Genetics Core Facility, and the Research Computing group at Harvard Medical School for assistance. We thank J. Neil (Walsh lab) for help with brain acquisition and IRB paperwork. We thank K. Stafstrom, W. Bainter, M. Reigle, and S. Weeks for assistance with IonTorrent chips. We thank all members of the Walsh and Park labs for critical feedback, especially Y-N. Kim, A. Kriz, E. Chun, K. Chatzipli, J. Markowski, and D. Gulhan for critical feedback on figures and text; S. Ehmsen on the design of figures, and J. Song for assistance with layout. This work was supported by the Stuart H.Q. and Victoria Quan Fellowship in Neurobiology (SNK), NLM grant T15LM007092 (VVV), NCI grant F31CA264958 (VVV), NINDS grant R01NS032457 (CAW), NIMH grant U01MH106883 through the Brain Somatic Mosaic Network (PJP, CAW), and the Templeton Foundation. C.A.W. is supported by the Allen Discovery Center program through The Paul G. Allen Frontiers Group. C.A.W is an Investigator of the Howard Hughes Medical Institute.

## Author Contributions

SNK, PJP, and CAW conceived the study. SNK designed and performed the experiments, with assistance from RND, SB, SK, RY, ML, LR, PPL, JG, RSH, RAS, and JWT. SNK and RND designed the targeted amplicon sequencing pipelines and led the corresponding bioinformatic analyses. SNK and RND designed and performed the MIPP-seq experiments and bioinformatic analyses. SNK, SB, PPL, SA, RAS, and AYH performed PRDD-seq experiments and analyses. VVV, YD, AB, and AYH led the bioinformatic analysis on the WGS dataset and WGS variant detection. VVV performed bioinformatic analysis for cell-lineage tree construction. SNK, SK, SA, and SB performed snRNA-seq experiments. SB and YD contributed additional snRNA-seq and snATAC-seq data. VVV performed snRNA-seq and snATAC-seq data analyses and cell-type analyses resulting from those datasets. AT and KMK constructed an automated Python/Perl workflow for high throughput data processing of targeted sequencing datasets and statistical analyses. MB performed additional statistical analyses. SNK, VVV, PJP, and CAW wrote the manuscript, assisted by RND and MB and with input from all authors. CAW directed the research with input from PJP.

## Methods

### Resource Availability

#### Lead Contact

Further information and requests for resources and reagents should be directed to and will be fulfilled by the lead contact, Christopher A. Walsh.

#### Materials Availability

This study did not generate new unique reagents.

#### Data and code availability

Code used for bioinformatic analysis described in Figures 1, 2, 5, and 6 and their associated supplemental content can be found at https://github.com/parklab/spatial_sampling_analysis. MosaicForecast can be found at https://github.com/parklab/MosaicForecast. Raw sequencing data from UMB4638 and UMB4643 is available from dbGaP under accession number phs001485.v2.p1 and from the NIMH Data Archive 111. Raw sequencing data from UMB5575 and UMB5580 and raw data from newly collected snRNA-seq from UMB4638 and UMB4643 will be made available through dbGaP. All mutations and their validation status, single-cell lineage information, and cell-type annotations will be made available through the Supplementary Materials, which will be released upon acceptance and publication. The code pertaining to the Excel/Python/Perl workflow to analyze MIPP-seq results and produce the content of Figures 3 and 4 can be found at https://github.com/soniankim/brain-clone-mosaic. Any additional information required to reanalyze the data reported in this paper is available from the lead contact upon request.

### Experimental Materials and Methods for variant discovery and validation

#### Processing of human tissues and DNA samples

Human tissues were obtained from the NIH NeuroBioBank at the University of Maryland Brain and Tissue Bank. Fresh-frozen post-mortem tissues from two neurologically normal individuals were used in this study: UMB4638 (a 15-year-old female) and UMB4643 (a 42-year-old female). UMB4638 died from motor vehicular injuries and UMB4643 died from cardiovascular disease. Both individuals had no known neurological or psychological diagnoses at the time of death. Both individuals were obtained as part of previous studies in our lab (Bizzotto et al., 2021; Lodato et al., 2015). All tissue samples were prepared according to standardized protocols (https://www.medschool.umaryland.edu/btbank/Researchers/Tissues-Collected and https://www.medschool.umaryland.edu/btbank/Medical-Examiners-and-Pathologists/Minimum-Protocol) under the supervision of the NIH NeuroBioBank ethics guidelines. Brodmann area identification and sampling were completed by the NIH NeuroBioBank at the University of Maryland Brain and Tissue Bank.

Cortical samples were biopsied from the left hemisphere of both individuals. For initial variant calling in both individuals, bulk samples were biopsied from the PFC and occipital lobe, specifically BA17 and BA18. Likewise, single cells for variant calling were isolated from PFC (UMB4638: coronal section 3; and UMB4643 coronal section 4). For downstream experiments, including validation experiments, related to these three areas, biopsies from BA9 (representing PFC), BA17, and BA18 were used.

Bulk DNA was extracted from tissues using the lysis buffer from the QIAamp DNA mini kit (Qiagen; Cat. 51304) with proteinase K digestion and RNase A treatment, followed by a phenol-chloroform extraction and alcohol precipitation.

Single nuclei were isolated by fluorescence-activated nuclear sorting (FANS) using an anti-NeuN antibody as a neuronal nuclei marker (Millipore, MAB377). Nuclei were lysed on ice in alkaline conditions, and whole-genome amplified using MDA, as previously described (Bizzotto et al., 2021; Lodato et al., 2015; Evrony et al., 2015; Dean et al., 2002).

#### Human sample preparation

We received tissue biopsies from two neurologically normal individuals, UMB4638 and UMB4643, from the NIH Neuro-BioBank. These samples have been used in prior publications for some limited variant discovery, WGS, and clonal analysis (Bizzotto et al., 2021; Lodato et al., 2015). The left hemisphere of each brain was analyzed, with the right hemisphere having been prepared by the NIH NeuroBioBank for histological analysis and thus unavailable for DNA sampling. Because these brains represent shared resources, many cortical regions had already been extensively sampled, especially primary motor cortex, primary somatosensory cortex, hippocampus, and other regions, and thus were unavailable for our analysis. These unavailable regions are indicated on the cortical maps in areas with no color shading and outlined with gray dashed lines outlining the representative BA regions and their unavailability.

The left hemispheres of each sample were sectioned coronally at ≈ 1 cm, according to neuropathological conventions and standard operating procedures. Approximate coronal section thicknesses were measured for coronal sections available for sampling (**Fig. S3A**). Samples were requested from all cerebral cortical BA regions available and identifiable, and from most if not all subcortical and non-cortical brain sites as well. Biopsies of cortical areas were cut by dieners at the NIH Neuro-BioBank, using extensive photographic maps and atlases of the human brain, recording the position of the sample relative to gyral landmarks, and the section number. Assignment of samples to BA regions comes from this biopsy process. Furthermore, photographs of coronal sections and tissues were taken before and after dissection for lucida tracing of biopsy locations within the coronal sections. Tissue samples are stored at -80°C until sample preparation. Sample preparations for bulk DNA extraction are as described before (Lodato et al., 2015; Bizzotto et al., 2021). Biological duplicates for BA9, BA18, and BA17 were isolated from the same tissue biopsy with the same protocol but separately prepared.

#### Library preparation and WGS for variant calling

Deep WGS (210X) on bulk tissue DNA was prepared using the Illumina TruSeq PCR-free preparation kit for paired-end barcoded WGS libraries. Paired-end sequencing (150 bp x 2) was performed on an Illumina HiSeq X10 (UMB4638 and UMB4643) or NovaSeq6000 (UMB5575 and UMB5580) instrument (Psomagen, Inc., Rockville, MD).

As described previously (Bizzotto et al., 2021), single neuronal nuclei were isolated using FANS with NeuN staining, a neuronal nuclei marker. Single neuronal sequencing was prepared by shearing 100 ng of DNA of each sample on a Covaris Ultra-Sonicator to yield ≈ 350 bp fragments. Paired-end barcoded WGS libraries were prepared using the Illumina TruSeq Nano LT sample preparation kit, and paired-end sequencing (150 bp x 2) was performed on an Illumina HiSeq X10 instrument. Library preparation and sequencing were completed at the New York Genome Center (New York, NY). Sequencing data of ten single prefrontal cortex neurons from each brain, which were selected based on low allelic and locus dropout rates, were made available from a previous study (Bizzotto et al., 2021).

#### Targeted amplicon sequencing of bulk DNA

In UMB4638 and UMB4643, we validated identified mutations using deep amplicon sequencing in 37 brain samples and 18 non-brain tissues samples (**Table S1.1**). Candidate sSNVs were selected based on parameters set by the amplicon design pipeline requiring primers mapping to unique genomic regions. Targeted regions were captured by the amplicon panel in bulk unamplified DNA samples from both brain and non-brain tissues (**Table S1.1**). Targeted sequencing of bulk DNA samples was completed using a custom designed amplicon pool and a custom library preparation and barcoding protocol. A custom amplicon panel for each individual was designed to target specific candidate sites using the Ion AmpliSeq Designer tool (Thermo Fisher Scientific). Each amplicon pair was designed to be unique and specific to a target candidate site. The amplicon panel was used in the initial targeted capture step with minimal PCR cycles to reduce artifacts from PCR amplification. The initial input amount of DNA was 20 ng per reaction, consisting of 9 μL of 2X custom AmpliSeq Primer Pool and 10 μL of 2X Phusion U Mastermix (Thermo Fisher Scientific, F-562). Targeted amplicon sequencing of bulk tissue DNA was prepared using a custom library prep protocol for paired-end barcoded WGS libraries. Paired-end sequencing (150 bp x 2) was performed on an Illumina HiSeq X instrument. Library sequencing was completed by Psomagen, Inc. (Rockville, MD).

For all targeted captures using the custom amplicon panel, two biological duplicate bulk DNA samples representing two separate extractions from the same tissue region were used. For technical replicates, each biological duplicate was prepared three times, for a maximum of 6 samples for each evaluated tissue. For controls, preparations were also performed using 1) nuclease-free water, 2) an unrelated male fibroblast genomic DNA sample (Promega, G1471), and 3) DNA from the other individual (i.e., using the custom panel specific to UMB4643 on UMB4638 bulk DNA). Specifically, regarding the region-validation experiments, the following samples for BA17 and BA18 were used: bulk tissue DNA samples used for the original sSNV detection, and a biological duplicate sample extracted similarly from the same tissue but not used for candidate sSNV discovery. For PFC, two biological duplicate samples were extracted similarly from within the same BA9 region; this exact BA9 tissue biopsy was not used for the original SNV detection. Custom amplicon panels were also used to target sSNVs in additional bulk DNA samples extracted from both brain and non-brain tissues.

### Bioinformatic methods for variant calling

#### Definitions

Mosaic fractions (MFs) are defined as twice the alternate allele fraction (2 x AAF), expressed as an average if multiple amplicons in MIPP-seq were designed to target the sSNV. Following convention, we define mild and extreme outliers as observations that are respectively 1.5-3 and at least 3 interquartile ranges (IQRs) beyond the upper (q1) and lower (q3) quartile values. For reference, the IQR is measured as the difference between the lower and upper quartiles. Mathematically, mild outliers are observations (x) that satisfy *x* < *q*1 − [1.5, 3] × *IQR* or *x* > *q*3 + [1.5, 3] × *IQR*, whereas extreme outliers satisfy *x* < *q*1 − 3 × *IQR* or *x* > *q*3 + 3 × *IQR*, where *IQR* = *q*3 − *q*1.

#### Statistical analysis

Statistical analysis, including counts, averages, and statistical tests are reported in figures and tables. Statistical analyses were performed in Microsoft Excel, Python, and R.

#### Whole-genome sequencing data processing and filtering

WGS reads generated from single neuronal sequencing were processed as previously described (Bizzotto et al., 2021). WGS reads from bulk tissue sequencing were prepared, in brief, by mapping reads on to the human reference genome (GRCh37) by Burrows-Wheeler Aligner (BWA) with default parameters. Duplicate reads were marked with MarkDuplicate of Picard tools (Broad Institute, 2019) and further post-processing was completed with local-realignment around indels and base-quality score recalibration using Genome Analysis Toolkit (GATK, version 3.5) (McKenna et al., 2010).

#### Somatic single nucleotide variant calling

sSNVs in bulk and single cell DNA WGS was called using MuTect2 (bulk calling) (Cibulskis et al., 2013), MosaicForecast (single-cell and bulk calling) (Dou et al., 2020), single-cell Mosaic Hunter (single-cell and bulk calling) (Huang et al., 2020, 2017), and a GATK-based triple-calling strategy (single-cell calling) (Lodato et al., 2015), as previously described. sSNV calling in bulk DNA WGS using MuTect2 (version *nightly*− 2016− 04− 25− *g*7*a*7*b*7*cd*) utilized the panel of normal tissue approach to complete a tissue-region versus tissue-region comparison to isolate for region-specific calls, promising greater sensitivity in detecting regional mutations. All calls (pass and non-pass) from MuTect2 were considered in order to increase sensitivity and minimize the loss of potentially rare sSNVs. All other sSNV detection parameters, along with the associated filter thresholds, for the other call-sets were as previously described (Bizzotto et al., 2021; Lodato et al., 2015; Dou et al., 2020; Evrony et al., 2015). The total sSNVs derived from all calling methods were then filtered for somatic mutations unique to each individual by excluding variants shared between both individuals. Variants located in segmental duplications and repetitive regions were also filtered out before designing the amplicon panel for targeted re-genotyping.

#### Estimating the ratio of the number of variants detected in one region versus another

Comparisons of the number of mosaic variants found in one brain region must take into account several technical factors. Mosaic variant discovery pipelines can exhibit different sensitivities based on the AAF of desired variants; for example, the sensitivity to detect mosaic variants at AAFs of <5% is significantly lower than for variants at AAFs in the 5-30% AAF range (Bizzotto et al., 2021). Higher-AAF variants can be shared across multiple brain regions, and brains can vary in the total number of variants discovered due to batch effects or sequencing platforms. Thus, we sought to estimate the ratio of variants present in BA17 versus BA8 (and similarly for BA9 versus BA17 or BA18), as a ratio would summarize a fundamental difference in the variants found in one region versus another due to one region having more region-specific variants or having variants at significantly higher AAFs. If two regions share a similar number and set of variants, then we assume that a variant detected in one of those regions would be found at a similar AAF in the other region. This latter case serves as our null model against which we can use to test whether the estimated ratio of variants in one region over another is significant.

In each of seven different AAF bins (1-2%, 2-3%, 3-4%, 4-5%, 5-10%, 10-20%, and 20-50%), we simulated the minimum number of sequencing reads at which we would detect a variant in the bin based on the AAFs of all variants found in this bin (our “threshold”). For each variant in each region, we simulated the number of sequencing reads at which it would be detected in that region as a binomial random variable with *N* equal to 250 reads and p equal to the AAF at which the variant is found in the region, and we retained the variant if the simulated number of reads exceeds that of our threshold. We estimated the projected number of similar variants that can be detected at the observed variant’s AAF by computing the reciprocal of the sensitivity estimated from a smoothed spline fitted to sensitivity estimates previously published for MosaicForecast (Bizzotto et al., 2021). To obtain the ratio of projected variants in that region, we summed up the numbers of projected variants; this sum is used to compute the ratio of variants that exist in one region versus another. The ratio under the null model is estimated with a similar procedure, except a particular variant’s AAF in a region is simply the mean of the AAFs across all tested regions. For example, a variant detected at AAFs of 2% in BA17 and 1% in BA18 in the same brain would be simulated at an AAF of 1.5% for the control. Simulations were conducted over 1000 iterations, and the 99% confidence interval was computed to provide an interval estimate of the ratio

#### Analysis of amplicon sequencing data

Sequencing reads were prepared by first trimming the reads for quality and removing any leftover adapter sequences from the reads using CutAdapt (Martin, 2011) (− *q* 20,−*u*−5, −*U*−5,−*a AGATCGGAAGAGC A AGATCGGAAGAGC*). Next, common sequencing artifacts were corrected using the Pollux software (Marinier et al., 2015) to generate both error-corrected and original fastq files using the following settings: −*p* −*n true*− *d true* −*h true*− *s f alse*− *f f alse*.

Reads were then mapped onto the human reference genome (GRCh37) by BWA-mem with default parameters. Further post-processing was completed with local-realignment around indels using Genome Analysis Toolkit (McKenna et al., 2010)] (GATK, version − 3.7, *T IndelRealigner*− − *filter*_*b*_*ases*_*n*_*ot*_*s*_*tored greedy*− 1200 *maxReads* 2000000 − *maxInMemory*1500000), using all InDels from gnomAD version 1 (Karczewski et al., 2020) as a control set. Finally, all primer binding sites were clipped from the sequencing reads using Bamclipper 93 and a bed file of all primers. Variants located within each amplicon were called using samtools mpileup version 1.3.1 ( − − *output*− *tags INFO*/*AD, DP, AD*− *Q* 20 − *q* 20) (Li, 2011). SNV variants were called alternate or reference using samtools mpileup (Li, 2011). Finally, VCFs for each sample were processed to include 50 nucleotides flanking each side of the targeted mutation for estimating the background error rates.

All variant calls were further validated to distinguish TPs from FPs and germline events using a combination of public databases (gnomAD version 3.92), manual review of genome mappability, control tissue sequencing, and the comparison of the original tissue in which a mutation was identified against other tissues in the individual. True positive mutations were defined as being high quality sites with good mapping, rare/absent in gnomAD version 3.92, absent in control DNA samples, alternate allele fraction (AAF) of 0.5% - 35%, and an allele depth (AD) > 2. Furthermore, the high confidence mosaic alleles were required to be detected within the tissues where they were originally identified. However, given some variability in amplicon sequencing depths, mutations not detected in the original tissue can also be considered as valid mutations given that they meet all other criteria and are present in multiple library preparations. Mutations additionally identified with an AAF suggestive of a germline event in control DNA samples (Promega genomic control and unrelated individual’s brain tissue), that appear as common high-quality germline events in gnomAD version 3 (Karczewski et al., 2020) with good read mapping, were flagged as true germline events. Alleles present in the control tissues, regardless of AAF, that are present in gnomAD version 3 (Karczewski et al., 2020), with poor quality flags and poor read mapping were further manually curated to confirm their FP status. Finally, any mutation consistently identified as a germline event across all tissues of a given individual, with an average AAF of 40-60% across all samples, were manually reviewed and classified as a true germline event.

#### Figures for lucida tracings and brain maps with Brodmann area annotations

Lucida tracings of sampled cortical sections (Figure S3A) were traced from photographs taken by the NIH NeuroBioBank at the time of tissue biopsy. Dashed lines indicate regions not present in photographs due to sampling prior to this study. Anatomy was extrapolated from records of sample locations, adjacent sections, photographs of right hemisphere formalin-fixed coronal sections, and atlases and MRI records of neurologically normal brain anatomy.

Lateral and medial cortical brain maps with Brodmann area annotations were adapted from the Brodmann (1909) areas (annotated) scene files for the left cortical hemisphere from the Brain Analysis Library of Spatial maps and Atlases (BALSA) database (Van Essen et al., 2017). Areas that are filled with color represent the corresponding MF of the sSNV in that BA sample.

### Generation and analysis of panel single-cell multiple displacement amplification (pscMDA) data

#### Alignment and genotyping of cells from pscMDA

We used multiple displacement amplification (MDA) to capture 122 brain-restricted sSNVs (56 for UMB4638 and 66 for UMB4643) across 1131 single nucleus genomes (563 in UMB4638 and 568 in UMB4643) taken from BA17, BA18, and BA9. We used cutadapt 90 with error rate set to 50% to aggressively trim adapters, partial adapter sequences, poly-G sequences, and polyX-sequences from demultiplexed FASTQ files of the pscMDA experiment. We aligned all reads to hg19 (GRCh37) using bwa-mem. We genotyped each mutation in our panel from the FASTQ data using procedures described before 16. Briefly, the genotyping model assumes that the posterior probability of a site being somatic-mutant in a cell can be computed from a binomial likelihood of observing alternative-allele backing reads at observed counts at probability p, i.e., the expected read fraction of a somatic-alt variant in the cell (ideally at 0.5 but potentially different given allele imbalances introduced during amplification). The posterior probability of a site being non-mutant at a site is also computed from a binomial likelihood of observing erroneous (i.e non-reference) reads at the site. The prior probability of a site’s genotype within a given cell is proportional to the observed read fraction of the mutant allele across all cells.

All parameters are estimated from heterozygous SNPs introduced in the panel and off-target amplifications that serve as examples of reference-homozygous sites. For instance, amplifications of UMB4643 sites in UMB4638 samples were used to estimate parameters for the reference-homozygous genotype in UMB4638. The two batches of sequencing data for the same cells and sites were genotyped separately before generating a consensus genotype matrix, using the estimated cell fraction of the variant to compute a binomial probability of the variant being somatic-mutant within a given cell across both batches. All parameter fittings were conducted using JAGS implemented through R, and code to genotype cells and sites is provided on the repository linked under the linked repository. We genotyped >85% of the sites across 1124 nuclei (Figure S2A, S2C), with cell fractions of each variant correlated closely with the MFs estimated from the AAFs measured by deep targeted sequencing of these variants (Figure S2B).

#### Construction of lineage trees

We assume that mutations evolve neutrally within the lineage (i.e., negligible chance of recurrent mutations newly arising on separate branches of a lineage tree without being inherited from a common ancestor). We applied scistree (Wu, 2020) to our consensus genotype matrix to filter out genotypes with this model (for example, for two sites [*A, B*], subpopulations exist with genotypes [0, 1], [1, 0], and [1, 1]) and impute compatible genotypes constrained by variants’ cell fractions. We used mpboot (Hoang et al., 2018) to construct maximum parsimony trees from this imputed genotype matrix. Genotype matrices from before and after imputation are reported in **Fig. S2A**.

#### Coalescent model for inferring variant origin timings and time elapsed during the observed lineage

We sought to infer the coalescent time of the entire population, which is the time elapsed between the population from which we have sampled our cells and its most recent common ancestor (MRCA). A population may develop from its MRCA depending on several parameters: mutation rate, fluctuations in population size over time, division rate (or birth and death rates), selection for specific mutations, and any other population structure (for example, migration or spatial segregation of the population). A phylogenetic tree represents the genealogical relationships of cells produced under these conditions, and a number of phylogenetic trees may be constructed to describe the different genealogies possible under the population parameters (given that we may end up observing a unique set of mutations or cells when sampling from the population). In any such tree, a node at which a tree branches off into descendant nodes is said to be the ancestral cell to those descendant cells, and the descendants are said to “coalesce” at this ancestor when we traverse up their branches to this ancestor. A mutation is mapped to a branch immediately leading up to a node if we find that cells descended from this node also carry the mutation. Under an assumed mutation rate, branch lengths can indicate the number of mutations mapped to them, which we assume is Poisson-distributed. We can use the number of mutations mapped to a series of branches connecting nodes to estimate the time elapsed between these nodes, each of which may represent an independent coalescence event. The coalescent time of the entire population is said to be the sum of all individual coalescent times in the lineage, or the sum of times between each coalescence event at which different portions of the population arrive at their common ancestor when traversing up the lineage tree.

All mathematical and algorithmic details for fitting the coalescent model are provided with the attached supplement (**Supplemental Methods**).

#### Calculating the regional restriction statistic

Coalescent timings were inferred and lineages were constructed agnostic to the regions where mutations were detected. To assess each clade’s association with different regions, we constructed the regional restriction statistic (RRS), which we formalized as the log-odds ratio of the distance between two cells in a clade belonging to the same region to the distance between cells from different regions. We envisioned this ratio as describing whether two cells from a particular clade and from the same region tend to be more closely related than if those cells were from different regions. For each pair of cells within each clade, we computed the ratio of the phylogenetic distance between cells within the same region to the distance between cells from different regions, and we estimated the mean and standard deviation of the distance ratios for each clade. The mean ratio is the RRS for a clade.

A RRS close to 1 suggests that cells within this clade are equally related whether in the same region or in different regions, suggesting that the variant’s dispersal throughout the cortex is not significantly shaped by or associated with regional separation. A RRS significantly above 1 suggests that cells in this clade are more related to clade-mates within the same region than across regions, suggesting that the clade is restricted mostly to one region or asymmetrically allocated to one region. A RRS significantly below 1 suggests that clades are more related to clade-mates from different regions than within the same region, suggesting that the clade widely populates other regions early on prior to the occurrence of later-stage variants. A null RRS can be defined using early-mosaic or germline mutations, which should precede the formation of brain regions and subsequent allocation of cells amongst them.

### Mosaic characterization using ultra-deep targeted sequencing (MIPP-seq)

### Preparation and capture of target sites using MIPP-seq

Mosaicism estimation of bulk tissue DNA was obtained using deep-targeted sequencing of regions captured by custom-designed primers, as described in a recently published method 36. When possible, ≥1 unique primers (termed “replicate primer pairs”) were designed to a SNV, with each additional replicate primer designed to stagger around the site of interest; this is to account for potential allelic dropout and imbalance, and to provide a more accurate mosaicism estimation of the targeted SNV. Every primer pair was designed with a sequencing adapter and unique barcode. Each primer pair was individually evaluated for a single expected product of correct fragment size and checked for efficiency. Custom-made multiplexed primer pools were generated and checked for primer cross-reactivity and capture efficiency. In brief, primer pairs were evaluated both independently and in pools, which were compared on a Tapestation D1000 ScreenTape system to check for proper product sizes. Replicate primer pairs targeting the same sites were placed in separate pools or used individually. Primer pairs that showed cross-reactivity with other primer pairs within a pool, such as abnormal fragment sizes, were isolated and ran in individual reactions as previously described 36.

The targeted sequencing was prepared by running a PCR with the primer pairs and 50 ng of bulk DNA input on low cycle number (20 cycles) (Doan et al., 2021). Libraries were prepared, and sequencing was performed on the Ion Torrent S5 sequencing platform. A calculation using an estimated 6-7 pg of DNA content per cell approximate ≈7,142-8,333 cells (Bä umer et al., 2018). Using this estimation, sSNVs with 0.1% MF, the lower limit of detection by this method, would represent ≈ 7-8 cells in the cellular population carrying the heterozygous sSNV.

#### MIPP-seq data processing, filtering, and analysis

Raw unmapped BAMs consisting of uniquely indexed amplicon sequences were converted to fastq using “bedtools bamtofastq” (Quinlan and Hall, 2010) prior to being demultiplexed into amplicon-specific fastq files based on their unique 15 nt barcodes with FASTX toolkit’s *f astx barcode splitter* (−− *bol*− − *mismatches* 3). Error correction was performed using Pollux (Marinier et al., 2015) ( −*n f alse d f alse*− *h true*− *s f alse f*− *f alse*), followed by barcode and quality trimming with CutAdapt (Martin, 2011) ( *u* 10 *q* 10). Each amplicon specific fastq was independently mapped against the human reference genome, hg19, using default settings in BWAmem. Local realignment was performed using GATK version 3.7 IndelRealigner (McKenna et al., 2010) ( − *greedy* 1200− *maxReads* 2000000 − *maxInMemory* 1500000) using high quality indels extracted from the gnomAD genomes database. Finally, primer binding sites were clipped using the bamclipper tool (Au et al., 2017) with default settings.

Each locus site in each amplicon was evaluated as carrying the alternate allele if it met the following criteria: 1) a minimum of 10,000 reads at the site of interest; 2) carrying the primary alternate allele called during initial variant discovery; and 3) the mosaicism at the given site is ≥ 0.1% MF (0.05% AAF). AAF averages were reported for those SNV sites with multiple (replicate) primer pairs. As previously described, the lower detection limit of mosaicism estimation using 50 ng of DNA input is 0.1% mosaicism (0.05% AAF) (Lodato et al., 2015). For the graphical presentation of mosaicism on the brain map figures, any sites with <0.1% mosaicism or carrying the reference allele, but passed the minimum total read limit (10,000 reads per site), were categorized as “alternate allele absent” for that tissue (represented as the shaded gray areas on the brain maps). If a given site yielded <10,000 reads at the site of interest, it was designated as inconclusive (represented as non-shaded areas with gray dashed lines on the brain maps).

Background error rates (**Fig. S3C**) were calculated as previously described (Lodato et al., 2015). In brief, background error rates of mutations were calculated using the average allelic fractions within 100 bases surrounding the targeted SNV in each amplicon. This represents the likelihood of generating a mutational artifact. If multiple (replicate) primer pairs were designed to the target site, then an average background error rate was calculated for that specific SNV across the relevant primer pairs.

All sSNVs followed for subsequent spatial mosaic analysis had background error rates below the lower technical limit for signal detection (0.1% MF), indicating the level of sensitivity provided by MIPP-seq (**Table S3.2**). A comparison of all sSNVs in biological duplicates of the same cortical areas shows that most sSNVs have similar MF values across biological replicates (**Table S3.3, Fig. S3D**). In all, 32 sSNVs and 27 sSNVs were studied for UMB4638 and UMB4643, respectively.

#### Grouping of SNVs into somatic and germline categories

SNVs were grouped into somatic (ultra-low mosaic, low mosaic, and higher mosaic) or germline categories based on the average mosaicism (2 × alternate allele fraction (AAF)) percent of a SNV across all evaluated samples. Grouping was also further confirmed by the mutation categorization completed by targeted amplicon sequencing (see **Table S1.2**). SNVs with an average AAF of ≥45% to 50% for a heterozygous SNV were grouped as germline mutations. Somatic mutations were categorized as higher mosaic sSNVs if the average mosaicism was between 10-90% MF. Low mosaic sSNVs carry a mosaicism of 2-10%. This category range is based on the lower limit of detection (10% MF) for standard sequencing technologies for mosaic mutations, including Sanger sequencing, pyrosequencing, and standard exome sequencing. Ultra-low mosaic sSNVs are sSNVs with an average mosaicism of ≤ 2% across all evaluated tissues. A previous study demonstrated the appearance of general restrictions within the cortex beginning at 4.3% mosaicism for heterozygous SNVs isolated from BA9, with mutations at >5% mosaicism appearing widely outside the brain (Bizzotto et al., 2021). An additional study evaluating early human development using early-occurring sSNVs showed that brain-specific progenitors produced clones with average MFs of <2.5% across the cortex in one individual (Dehay et al., 1993).

#### Analyses of MIPP-seq data across multiple cortical regions

Starting with an *m × n* MP matrix containing *m* mosaics and *n* tissues, we used the function *get* − *summary*− *stats*() in the R-package “rstatix” to compute summary statistics for each mosaic, which include interquartile ranges (IQRs), and upper (q1) and lower (q3) quartile values. For reference, the IQR is measured as the difference between the q3 and q1. The outliers for each tissue were identified using the *identi f y outliers*() function in the R-package “rstatix”. Following convention, we define mild and extreme outliers as observations that are respectively 1.5-3 and at least 3 IQR beyond the upper (q1) and lower (q3) quartile values. Mathematically, mild outliers are observations (x) that satisfy *x* < *q*1− [1.5, 3] × *IQR* or *x* > *q*3 + [1.5, 3] × *IQR*, whereas extreme outliers satisfy *x* < *q*1− 3 × *IQR* or *x* > *q*3 + 3 × *IQR*, where *IQR* = *q*3− *q*1. Because we consider the mosaic data to be paired across tissues, we considered using a 1-way repeated measures analysis of variance (ANOVA) using the functions *aov*() package “stats”. The four assumptions for ANOVA include 1) independence of observations, 2) no significant outliers, 3) normality, and 4) homogeneity of variances. Due to the tissue sampling methods and nature of SNVs, the observations were considered independent. Outliers were identified as above. Normality was verified via QQ plots (using function *ggqqplot*() in R-packages “ggpubr” and “ggplot2”) and the Shapiro-Wilk test using the *shapiro test*() function in the R-package “rstatix”. Homogeneity of variances was verified (not shown in data) using the *levene test*() function in the R-package “rstatix”. As the above assumptions for ANOVA were violated, it was necessary to perform a non-parametric analysis of variance with Friedman’s test, by using the *f riedman test*() function in the R-package “rstatix”.

Post hoc analysis was conducted for all tissue pairs, including biological replicates in BA9, BA18, and BA17. The *shapiro test*() function was once again used to determine if the difference in mosaic fractions between tissue pairs was normally distributed, to determine whether to use the t.test() function (R-package “stats”) for the paired t-test (if normally distributed) or the *wilcox*.*test*() function (R-package “stats”) for the Wilcoxon Signed Rank test (if not normally distributed). The Benjamini-Hochberg procedure was performed to reduce the false discovery rate (FDR) in the multiple comparisons, by using the *p*.*ad just*() function in the R-package “stats”.

The correlation of mosaics between tissues was determined using the *cor*() function in R-package “stats”. However, the paired nature of the mosaic fractions between tissues, and the wide variability of mosaic fraction values between sets of mosaics, resulted in spuriously high correlation coefficients. Future work will require normalization of data to account for the variability of mosaic fraction prior to performing correlation.

Additional R packages, UpsetR (Conway et al., 2017) and core Tidyverse packages (Wickham et al., 2019), were used for figure generation.

### Analysis of cell types using single-nucleus (sn)RNA-seq and snATAC-seq data

#### snRNA-seq and snATAc-seq processing and analysis

snRNA-seq 10X Chromium Genomics datasets (Zheng et al., 2017) were prepared from three different cortical areas from UMB4638 and UMB4643. 10X single nuclei RNA sequencing data was generated from sorted cells using either DAPI or NeuN (neuronal nuclei marker) from the BA17 and BA18 areas from both individuals. Previously published DAPI-sorted snRNA-seq libraries from BA9 were also analyzed from both individuals (Bizzotto et al., 2021). In total, 72,839 cells were analyzed between both individuals.

We processed snRNA-seq data through CellRanger version 4.0.0 from the 10X website, and we aligned cells to an hg19 ENSEMBL version 17 reference. We ran cellranger count for each sample to produce BAM files and per-sample features, barcodes, and counts matrices and cellranger aggregate to produce a combined set of matrices. We ran Garnett (Pliner et al., 2019) on default settings to annotate cell types in our aggregated genes-to-cells counts matrix, using markers for 16 different neuronal, glial, and non-brain-cell types downloaded from the Allen Brain Atlas (Hodge et al., 2019). We used label transfer to annotate the snATAC-seq data based on snRNA-seq annotations using Seurat (Stuart et al., 2019).

#### Inference of snRNA-seq clades and excitatory/*inhibitory populations*

In snRNA-seq data, at the sites of sSNVs with alternative alleles known from our WGS experiments, each cell reported average coverage of 1-4 reads per site in snRNA-seq (maximum 64 reads per site) and between 1-2 reads supporting the somatic alternative allele (maximum of 16; **Fig. S5A-B**). Between 4-27% of cells tagged at least one somatic variant’s alternative allele, and≈ 10-20% of all cells sequenced per cell type reported coverage at a somatic alternative allele (**Fig. S5D**).

We grouped single cells from our snRNA-seq and snATAC-seq datasets based on shared, identifiable somatic variants. We constructed a genotype matrix for 12,381 total single cells (8,279 in UMB4638 and 4,102 in UMB4643) across both brains’ snRNA-seq and snATAC-seq datasets and applied Louvain clustering to identify 82 groups in UMB4638 and 83 in UMB4643, with cells in cluster sharing one or more common variants (**Fig. S5F**). We estimated the cell type composition of each cluster to identify patterns using annotations provided by Garnett.

Given the coverage of somatic mutations in our single-cell transcriptomic and chromatin accessibility datasets, we sought to identify clusters of cells that share common sets of variants and identify their cell type compositions. We constructed an adjacency matrix that reports the number of variants shared by each pair of cells, and we applied Louvain clustering to identify groups of cells that share common variants. Each Louvain cluster represents a set of cells that shares a common set of variants and any other variants that might be represented exclusively within a subset of those cells.

Garnett’s annotations were used to mark the compositions of cell types within individual Louvain clusters. However, due to technical constraints on single-cell sequencing, significant variation exists in the number of cells within each cluster (between 21-210) with a significant number of clusters consisting of only a single cell. Thus, the estimated percentage of a cell type within a small cluster would be more prone to fluctuations in the cluster size than would an estimate for a large cluster. We employed empirical Bayes methods to generate an estimate of cell type compositions while controlling for cluster size and the number of variants represented in the cluster (Robinson, 2017). We focused on the proportions of excitatory and inhibitory neurons. We modeled the number of cells in each cluster of size *N* coming from a cell type as a beta-binomial random variable, in which the observed number of cells *X* depends upon parameters μ and σ. For each cell type, we used beta-binomial regression through the “aod” package to regress the *X* and *N* − *X* on the number of variants represented in the cluster and the log10 cluster size. This regression model yielded estimates of μ_0_ for each cluster and a shared σ_0_, both of which were used to generate a prior distribution of cell type composition for each cluster. The posterior estimate of cell type composition was derived by computing

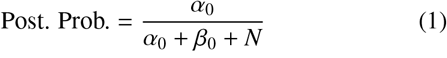

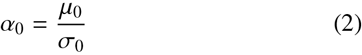

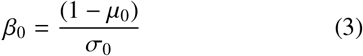

### Lineage and cell-type annotation using PRDD-seq

Lineage clading and cell-type analyses of UMB4638 was completed using PRDD-seq, as previously described (Huang et al., 2020). We performed additional cell-type analysis using sSNVs in UMB4638 and UMB4643 with PRDD-seq, completed as described (Huang et al., 2020). Designated marker genes used for cell-type and layer identification are described in (Huang et al., 2020).

### For Microsoft Excel and Python analyses

Analysis related to MF determination and background error rates are as previously described 17. Work was initially completed in Microsoft Excel, and later adapted to an automated Python/Perl workflow that processed data in a high throughput manner. Briefly, all sites were first checked to ensure they meet the minimum QC metrics described above (e.g., ¿10,000X depth, and detected alternate allele matches expected alternate allele from WGS). Next, the AAF at the variant position was extracted for each amplicon, with the average and 95% confidence interval being calculated for each variant using the 2+ independent amplicons, if applicable. Next, background error rates for each amplicon were measured as the average AAF across the flanking 100 nts proximal to the target variant, and averages and confidence intervals for error rates were further calculated across replicate amplicons. Finally, the average AAFs of the targeted variant were directly compared against the average background errors using a t-test. To ensure that error correction using Pollux did not introduce any errors in the data, assessments were performed using both the original (i.e., uncorrected) and error-corrected sequencing data.

