## Supplemental File for "Cell lineage analysis with somatic mutations reveals late divergence of neuronal cell types and cortical areas in human cerebral cortex"

### Main and Supplemental Tables

*S0.1.* ***Table 1: Mosaic values of variants defined by MIPP-seq and estimated mosaic fractions for sites and samples that were REF/ALT tagged from amplicon sequencing data***

Samples with no MIPP-seq data due to technical limitations were tagged as REF/ALT based on amplicon sequencing data. Samples tagged as REF from amplicon sequencing data were estimated to be reference allele carrying samples. Samples tagged as ALT from amplicon sequencing data received a mosaic estimation by averaging the MIPP-seq-measured mosaic values of its nearest topographic-neighboring tissue samples.

*S0.2.* ***Table S1, Related to Fig. 1***

- **Table S1.1**: List of sample locations and embryonic germ layer origin of human samples, Related Figure 1. All samples are from the fresh-frozen tissues of each individual. Coronal brain sections were biopsied from the left hemisphere. Cortical regions are listed from anterior to posterior. Additional brain and non-brain tissues are listed accordingly. Regions or tissues marked as NA are missing due to sampling prior to this study. Right-most column is the corresponding embryonic germ layer it was derived from during development. Adrenal tissue was not specified whether the sampled was biopsied from adrenal medulla (derived from ectoderm) or adrenal cortex (derived from mesoderm), so it is listed as mesoderm-ectoderm.
- **Table S1.2**: Number of amplicon-validated sSNVs that were rediscovered in the original region of variant discovery, Related to Figure 1.
- **Table S1.3**: Corresponding table of numerical counts to Figure 1E, Related to Figure 1. One indicates presence and zero indicates absence of sSNV in sampled tissue.
- **Table S1.4**: Variants called from UMB5575 and UMB5580
- **Table S1.5**: Validation of variants from UMB5575 and UMB5580. Validation was conducted by amplicon sequencing. Amplicons were submitted to Azenta Life Sciences for library preparation and sequencing using the Amplicon EZ protocol

*S0.3.* ***Table S2, Related to Fig. 2****: Tables of genotyping statistics from pscMDA of individual brains and batches*

- **Table S2.1-S2.8**: Genotype matrices for individual batches from each brain. Raw (from genotyping pipeline described in Methods) and imputed (from scistree as described in Methods) genotypes are presented.
- **Table S2.9-S2.14**: Genotype matrices from merged batches. Final genotypes, along with posterior probabilities of each cell being reference-homozygous or somatic-mutant are presented.
- **Table S2.15-S2.16**: Imputed genotypes from merged batches. Imputation was conducted using scistree (see Methods).
- **Table S2.17-S2.20**: Genotyping statistics for each brain and each batch

*S0.4.* ***Table S3, Related to Fig.* ?? *and Fig.* ??***: Tables of genotyping statistics from pscMDA of individual brains and batches*

- **Table S3.1**: Tables listing mosaic values from MIPP-seq measurements, Related to Figure 3. Minimum read depth: 5,000 reads per amplicon per site in order to confidently estimate mosaic fractions down to 0.1% mosaic fraction. Samples with no MIPP-seq data due to technical limitations were tagged as REF/ALT based on amplicon sequencing data.
- **Table S3.2**: Corresponding table of values to **Fig. S4A**.
- **Table S3.3**: Corresponding table of values to **Fig. S4B**.
- **Table S2.17-S2.20**: Genotyping statistics for each brain and each batch

*S0.5.* ***Table S4, Related to Fig. 5 and Fig. 6*** *: Results are presented for UMB4638 (S4.1, S4.3) and UMB4643 (S4.2, S4.4) for snRNA-seq (S4.1, S4.2) and snATAC-seq (S4.3, S4.4)*

*S0.6.* ***Table S5, Related to Fig.* ??**

- **Table S5.1**: PRDD-seq evaluated sSNVs for cell-lineage and cell-type integrated analysis on UMB 4638. List of clones for UMB 4638 evaluated using PRDD-seq. Three groups were identified (clades A, B, and C), and ordered in generation sequence with 1 being the earliest generated clone in that group (Huang et al., 2020). PRDD-seq related diagrams combined A2a and A2b for a combined A2 result, and A5a and A5b for a combined A5 result. PRDD-seq analysis completed in BA9/prefrontal cortex.
- **Table S5.2**: Number of cells genotyped and number of cells containing a positive genotype for each mutation assessed via PRDD-seq. Positive genotype indicates presence of alternate allele (left column).

### Supplemental Methods

#### *Fitting the coalescent model*

*Assumptions underlying the model*

The two individuals contributing UMB4638 and UMB4643 were neurotypical and had no apparent significant cortical malformations, and the mutations contributed by these individuals in our panel assumed to be neutral (i.e., no evidence for positive or negative selection at the loci where alleles were discovered). Thus, we modeled the lineage tree data as arising from a population that follows a Wright-Fisher model (Tran et al., 2013), in which we assume that genetic drift is the main contributor to somatic mutational variation across the cortex. We assumed the following:

1. The effective population size (i.e., the expected overall number of cells contributing to the mutations in the lineage) is constant across generations during the development of the lineage but otherwise unknown and to be inferred during model fitting.
2. Although the cell population expands in the development of the cortex, we assume that the effective population size harboring the observed number of variants is significantly smaller than the entire population size of the cortex at the time of development, which can help simplify computations.
3. As the average allele frequency of rare mosaic variants does not differ significantly across the three regions (based on **Fig. S2**), we can also assume that the mutation rate does not significantly differ across cortical regions. Region-dependent mutation rates would introduce significantly different numbers of alleles in each population and thus create significant differences in average mutation frequencies.
4. Finally, we assume that cell divisions are independent of one another and do not otherwise influence one another.

*Description of the model and the method used to fit parameters*

We adapted the standard Kingman coalescent model (Kingman, 1982) to conduct inference. Equations to estimate the coalescence times of the entire population and the individual depend upon a number of parameters:

1. The population size $N$.
2. The number of observed mutations. A subset of these mutations will vary across the cells, which are typically referred to as the number of segregating sites $S$.
3. The number of sampled cells $n$.
4. The mutation rate $\mu$.
5. A phylogenetic tree $\psi$ constructed using the observed mutations.

The time between two coalescence events $j$ and $j-1$, which we define as the coalescence time $T_j$, sees the sampled number of cells in our sample shrink from $n_j$ to $n_{j-1}$, where $n_j$ is the number of sampled cells still extant within the population at $j$. During this time, two cells in the population coalesce into their MRCA; for a large population, the probability that two cells coalesce with time $T_j$ can be modeled as an exponentially-distributed variable

$$T_j \sim Exp(\frac{(n_j(n_j-1))}{2}) \tag{S1}$$

$$E[T_j] = \frac{2}{(n_j(n_j-1))} \tag{S2}$$

where $E[T_j]$, the expected value of the coalescence time, is equivalent to the probability that the two cells are selected from the population to coalesce.

The estimated coalescence time of the entire subpopulation that we have identified is as follows.

$$T = \sum_{j=1}^{n} T_j \tag{S3}$$

$$\begin{aligned} E[T] &= \sum_{j=1}^{n} E[T_J] \\ &= \sum_{j=1}^{n} \frac{2}{(n_j(n_j-1))} \end{aligned} \tag{S4}$$

The expected value of the coalescence time, which is obtained by the laws of total expectation, represents a summary statistic of the average time for the sampled set of cells to coalesce into their MRCA.
The phylogenetic tree $\psi$ can be built from such a sampled population of $n$ cells with $S$ observed segregating sites, i.e., mutations whose genotypes differ amongst the cells and could therefore be mapped to the branches of the tree. If the mutation rate per cell is $\mu$, then $\theta = 2N\mu$ represents the population-wide mutation rate. Given that mutations accumulate along branches of $\psi$ (representing the accumulation of mutations in the lineage), the sum of branch lengths in $\psi$ is

$$L = \sum_{j=1}^{n} T_j \tag{S5}$$

The expected sum of branch lengths, $E[L]$, represents another summary statistic that we can infer about our sample.
Acceptable values of our parameters $\{E[T], E[L], \theta, N, \mu\}$ will yield a high likelihood for the observed value of $S$, and we seek to generate posterior distributions for our parameters to describe the timing of the lineage process. We used rejection sampling (Tavaré et al., 1997; Beaumont et al., 2002; Csilléry et al., 2010) to determine values of $\{E[T], E[L], \theta, N, \mu\}$ by computing the following in sequence:

$$\mu = 10^{-9} \tag{S6}$$

$$N \sim \frac{1}{\eta \log 10}, \ \eta \in [10^3, 10^9] \tag{S7}$$

$$\theta = 2N\mu \tag{S8}$$

$$E[T] = \sum_{j=1}^{n} \frac{2}{(j(j-1))} \tag{S9}$$

$$E[L] = \sum_{j=1}^{n} \frac{2}{((j-1))} \tag{S10}$$

$$s \sim Pois(\lambda), \ \lambda = \frac{1}{2}E[L]\theta \tag{S11}$$

The value of $\mu$ was chosen as the order-of-magnitude estimate for the somatic mutation rate as determined from past studies of the single-cell somatic mutation rate. The lower bound of the range of population sizes we sampled is the next-highest order of magnitude of our observed sample size in each brain (560 cells rounded up to 1000 cells), while the upper bound is one order of magnitude short of the approximately 85 billion cells within the human brain based on existing estimates (Herculano-Houzel, 2009). The acceptance probability for the parameters estimated above is

$$\text{Accept. Prob.} = \frac{Pr(X = s; \lambda)}{Pr(X = s; S)} \tag{S12}$$

where $s$ represents the sampled value of the number of segregating sites for a value of $\lambda$ based on the proposed sets of parameters. The null model assumes that the number of segregating sites in the population is the same as the observed number of segregating sites $S$ in the sample, which is obtained directly from the genotype matrix.
We conducted 5000 iterations of rejection sampling and retained parameters values with acceptance probability greater than 0.95. The accepted values of $\{E[T], E[L], \theta, N, \mu\}$ can be used to compute posterior estimates of these parameters to describe the properties of the population that gave rise to our sample and its lineage.

*Modifications for estimating the per-variant time-of-origin (TOO)*
A modified procedure was conducted to estimate the coalescence times of subpopulations of cells that all share the same variant and, by extension, the time at which the variant arose in the population. We assume that our phylogenetic tree $\psi$ contains a subtree $\psi_i$ for each mutation $i$ that was mapped to a branch and corresponds to the portion of the tree that descends from an ancestral branch to which mutation $i$ is mapped. We define $S_i$ as the number of segregating sites in $\psi_i$. Given the parameters that we estimated above, we now seek to estimate additional parameters $\{A_i, A_m, L_i, T_i, T_{i,m}\}$, where $A_m$ is the number of progenitors of the entire population, $A_i$ is the number of progenitors of the subpopulation, $T_i$ is the time for the subpopulation to coalesce to its MRCA, $L_i$ is the sum of the lengths of all branches of $\psi_i$, and $T_{i,m}$ is the time between the MRCA of the entire population and the MRCA of the subpopulation.

The values of $\{A_i, A_m, L_i, T_i, T_{i,m}\}$ are iteratively inferred until$A_i = 1$, i.e., when the subpopulation coalesces to its single MRCA. Using initial values of $A_i = n_i$, $A_m = n$, and $T_i = 0$,

$$W = \frac{2}{(A_m(A_m - 1))} \tag{S13}$$
$$T_i = T_i + W \tag{S14}$$
$$L_i = L_i + A_i W \tag{S15}$$
$$p = \frac{(A_i(A_i - 1))}{(A_m(A_m - 1))} \tag{S16}$$
$$A_m = A_m - 1 \tag{S17}$$
$$U \sim Bern(p) \tag{S18}$$
$$A_n = A_n - U \tag{S19}$$
$$T_{i,m} = \begin{cases} 0, & \text{if } A_m = 1 \\ \sum_{j=1}^{A_m} W_j, & W_j \sim Exp(\frac{j(j-1)}{2}) \end{cases} \tag{S20}$$
$$T_m = T_i + T_{i,m} \tag{S21}$$
$$\tag{S22}$$

Until $A_i \leq 1$, where $T_m$ is the coalescent time of the entire tree and $T_i$ is the coalescent time of the subtree with ancestral mutation $i$. Then, the acceptance probability of the parameter set $\{A_i, A_m, L_i, T_i, T_{i,m}\}$ is

$$\begin{aligned} \text{Accept. Prob.} &= \frac{Pr(X = S_i; \lambda)}{Pr(X = S_i; S_i)} \\ \lambda &= \frac{L_i \theta}{2}, X(\lambda) \end{aligned} \tag{S23}$$

$\theta$ is derived from the estimate of the whole-tree coalescent above, as we assume that the population mutation rate does not change significantly throughout the lineage. Rejection sampling is run for 1000 iterations for each variant. Each variant's $T_{i,m}$ is our estimate for the time when a variant is believed to have arisen after the start of the whole-tree lineage.

*Conversion of coalescent time parameters to real-world time estimates*

The estimated coalescent time parameters $\{T, T_i, T_{i,m}, T_m\}$ must be converted to real-world time units. To do so, we rely upon our previously-obtained estimate for $N$, and we assumed a division rate of 250,000 new cells per minute in the developing brain (Ackerman, 1992). Thus, the number of weeks within any estimate for coalescent time from our procedure is

$$\text{Number of weeks} = \frac{TN}{(\log_2 \frac{2.5\times10^5 \text{ new cells}}{60 \text{ seconds}} \times 8.64 \times 10^4 \frac{\text{seconds}}{\text{day}} \times 7 \frac{\text{days}}{\text{week}})} \tag{S24}$$

The $\log_2$ transformation allows for converting the number of new cells per second to the number of new cell divisions per second, as each coalescent event implicitly represents a new cell division event. $N$ is an estimate of the size of the effective population, whose members' divisions generate the sample of cells and mutations that we have observed.

The coalescent time of single-cell mutations in our tree is assigned to the total coalescent time of the entire tree as a "censored" estimate of the time-of-origin. The data does not allow us to rule out that the variant is present in additional cells in the underlying population from which we took our sample, but the coalescent time of the whole lineage does provide an estimate for the latest time at which single-cell mutations occurred.

The "grid" and "igraph" packages were used to construct the variant timelines, and custom code to produce the timeline plots in **Fig. 2** is provided in the linked code repository.

### Supplemental Figures

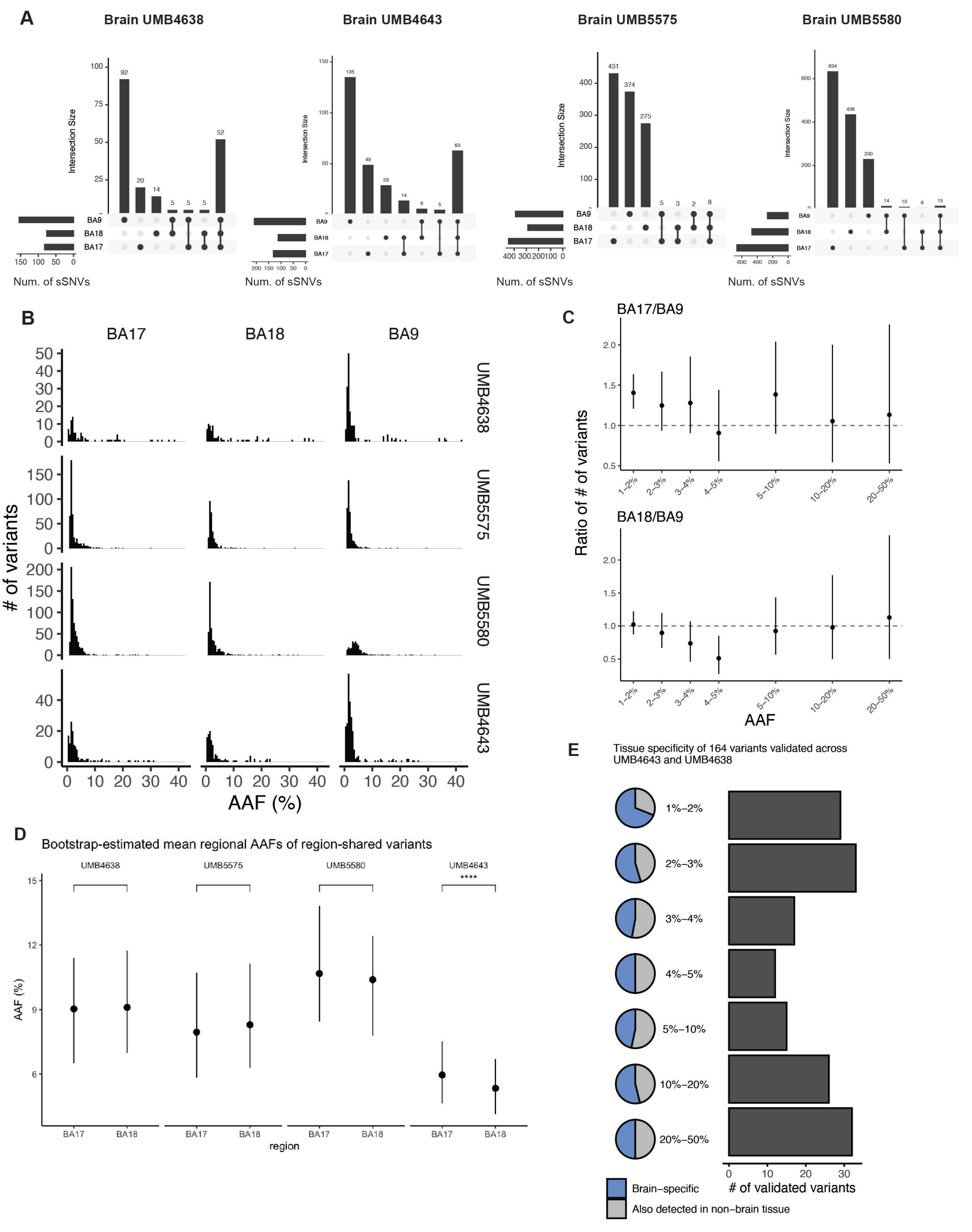


Figure S1: **Differences in sSNV counts across different cortical regions, Related to Fig. 1.** **(A)** Overall number of mutations called in >200X WGS from UMB 4638, UMB4643. UMB5575 and UMB 5580, and categorization of the distribution of each called variant. **(B)** Allele frequencies of sSNVs identified per brain and per cortical region. **(C)** Ratio of the sSNV count between BA17 and BA9 ("BA17/BA9"), and BA18 and BA9 ("BA18/BA9") for each alternate allele frequency range. BA9 showed higher sSNV counts than either BA17 or BA18, again at lower AAF. **(D)** Bootstrap estimates of the average AAFs of variants shared between BA17 and BA18. **(E)** Subset of amplicon-validated sSNVs present in tissues derived from major germ layers. Most validated sSNVs were found in at least 1 tissue derived from each embryonic germ layer, 44 were limited to ectoderm in UMB4638 and UMB4643, suggesting these sSNVs occurred after gastrulation.

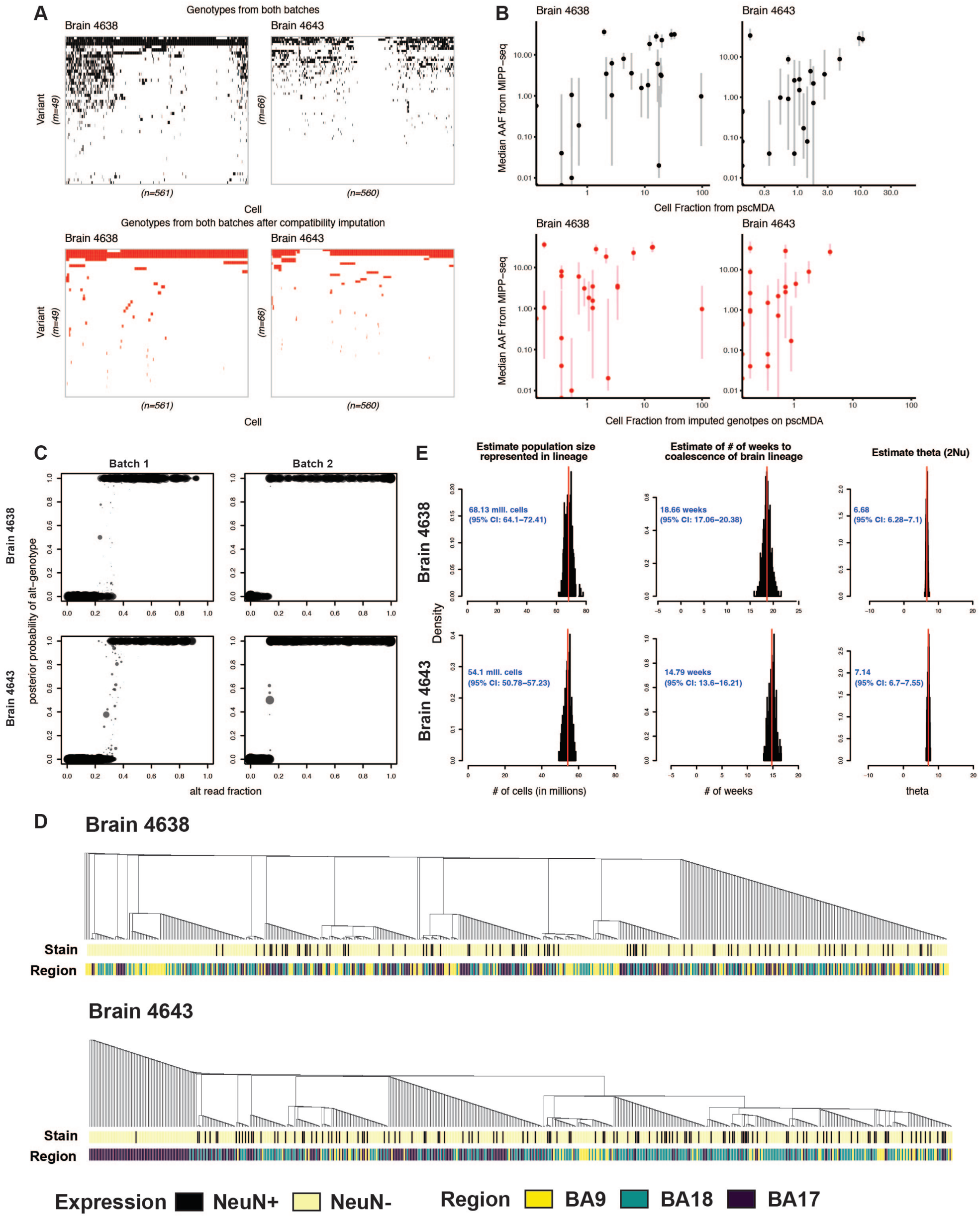


Figure S2: **Construction of single-cell lineage trees and inference of lineage parameters for UMB4638 and UMB4643. Related to Fig. 2.** **(A)** Genotype matrix of 561 (Brain 4638) and 560 (Brain 4643) single cells. Cells were sequenced using panel single-cell multiple displacement amplification (pscMDA) at select somatic mutation sites previously identified from 210X WGS and validated using IonTorrent PCR. *Top*: cells were sequenced across two batches, with each individual batch genotyped before integration of genotypes. Shown are the integrated genotype matrices. *Bottom*: scistree (Wu, 2020) was used to impute genotypes for single cells to ensure that all single cells' genotypes are compatible with the infinite sites assumption while ensuring that high-confidence genotypes are retained and that the phylogeny of the single cells is retained as much as possible. In each matrix, hierarchical clustering of the cells and variants was conducted independently of other matrices. **(B)** Comparisons of cell fractions inferred by pscMDA and MIPP-seq. Cell fractions inferred from the integrated genotype matrix (top, black) and the post-imputation genotype matrix (bottom, red) of pscMDA are plotted against the alternate allele fractions (AAFs) from MIPP-seq (IonTorrent-based sequencing of candidate somatic variants). For some variants, MIPP-seq variants are too low to confidently distinguish above error read fractions. Vertical bars (top, grey; bottom, pink) indicate the range of MIPP-seq AAFs across 21 brain regions and structures where the variant was sequenced in the corresponding individual, with points' y-values reporting the median across tissues. **(C)** Posterior probability of cells carrying the somatic-alt allele at each site. Each point represents a site within a specific cell. Points are sized by the log10 total coverage (with the log-value divided by 100 to determine point size). **(D)** Phylogenetic trees constructed from pscMDA cells. Branches are colored by the variant mapped to them. The heatmaps below the tree mark the nucleus stain (NeuN+/-) and region (BA17, BA18, BA9) from which the variant originates. **(E)** Posterior distributions of whole-lineage coalescent parameters. See **Methods** for details on model specification and fitting.

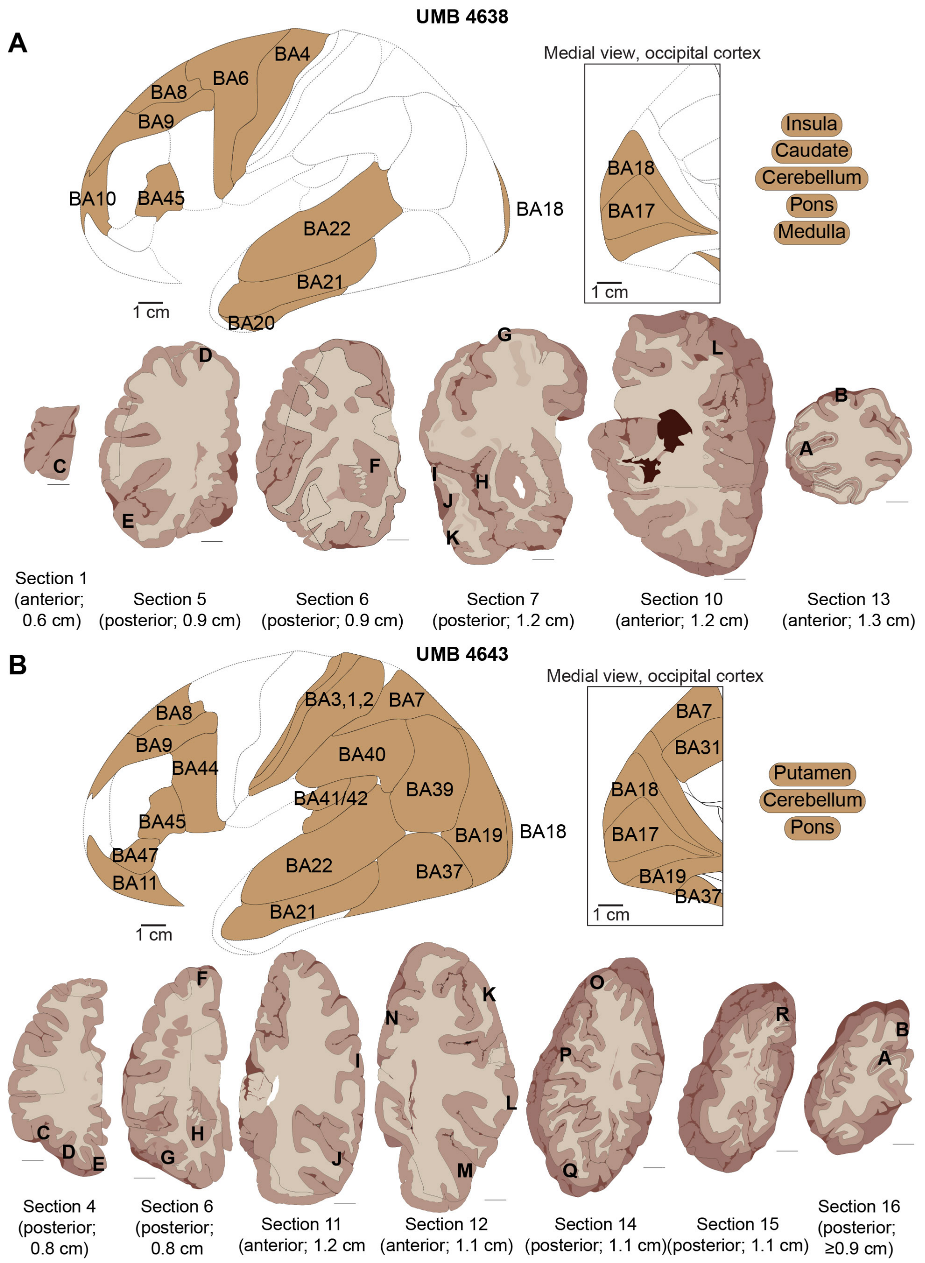


Figure S3: Continued on the following page.

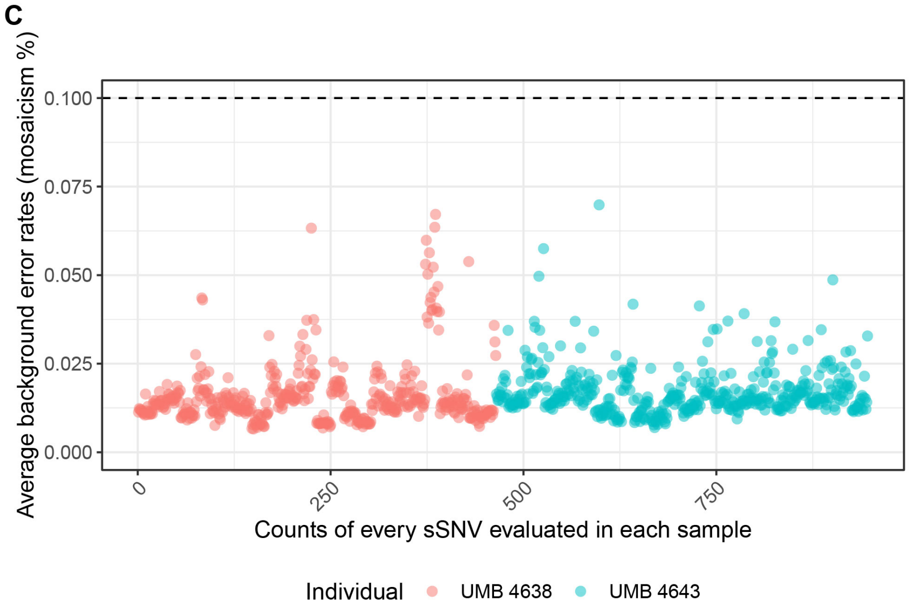


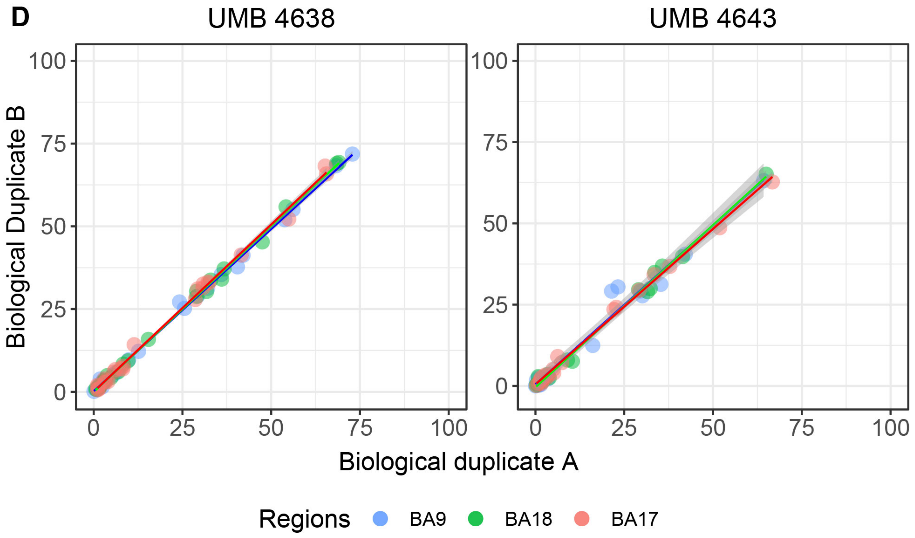


Figure S3: **(A and B)** Locations of cortical, subcortical, and non-cortical brain samples used for mosaic characterization and distribution analysis. Locations of samples from UMB4638 (A) and UMB4643 (B). For each panel, the top half shows the cortical brain map of the left hemisphere with BA annotations (see methods) indicate which BAs are represented in the study. Shaded regions indicate the BA represented by the obtained tissue biopsy. Some regions were unavailable due to prior sampling of these brains; these are indicated by the non-shaded regions with dashed grey lines. The bottom half shows lucida tracings of sampled cortical sections (see methods) indicate the BA sampled from that coronal section. Viewing orientation and the thickness of the coronal section are listed below each section. All sections are scaled; scale bar is 1 cm. (A) Sampled tissues (in UMB4638) are listed as: (C) BA10; (D) BA8; (E) BA45; (F) Caudate nucleus; (G) BA6; (H) Insula; (I) BA22; (J) BA21; (K) BA20; (L) BA4; (B) BA18; and (A) BA17. BA9 (not marked) is from section 5. Not shown here are additional brain tissues: pons, medulla, and cerebellum. (B) Sampled tissues (in UMB4643) are listed as: (C) BA45; (D) BA47; (E) BA11; (F) BA8; (G) BA44; (H) Putamen; (I) BA40; (J) BA41/42; (K) BA3/1/2; (L) BA22; (M) BA21; (N) BA31; (O) BA7; (P) BA39; (Q) BA37; (R) BA19; (B) BA18; and (A) BA17. BA9 (not marked) is from section 6. Not shown here are additional brain tissues: pons and cerebellum. (C) Average background error rate for sSNVs in each cortical region using MIPP-seq for UMB4638 and UMB4643. (D) Correlation of MIPP-seq mosaic fraction estimate across two biological duplicates (A and B; not related to the "A" and "B" clades assessed in **Fig. 3**) taken for each cortical region.

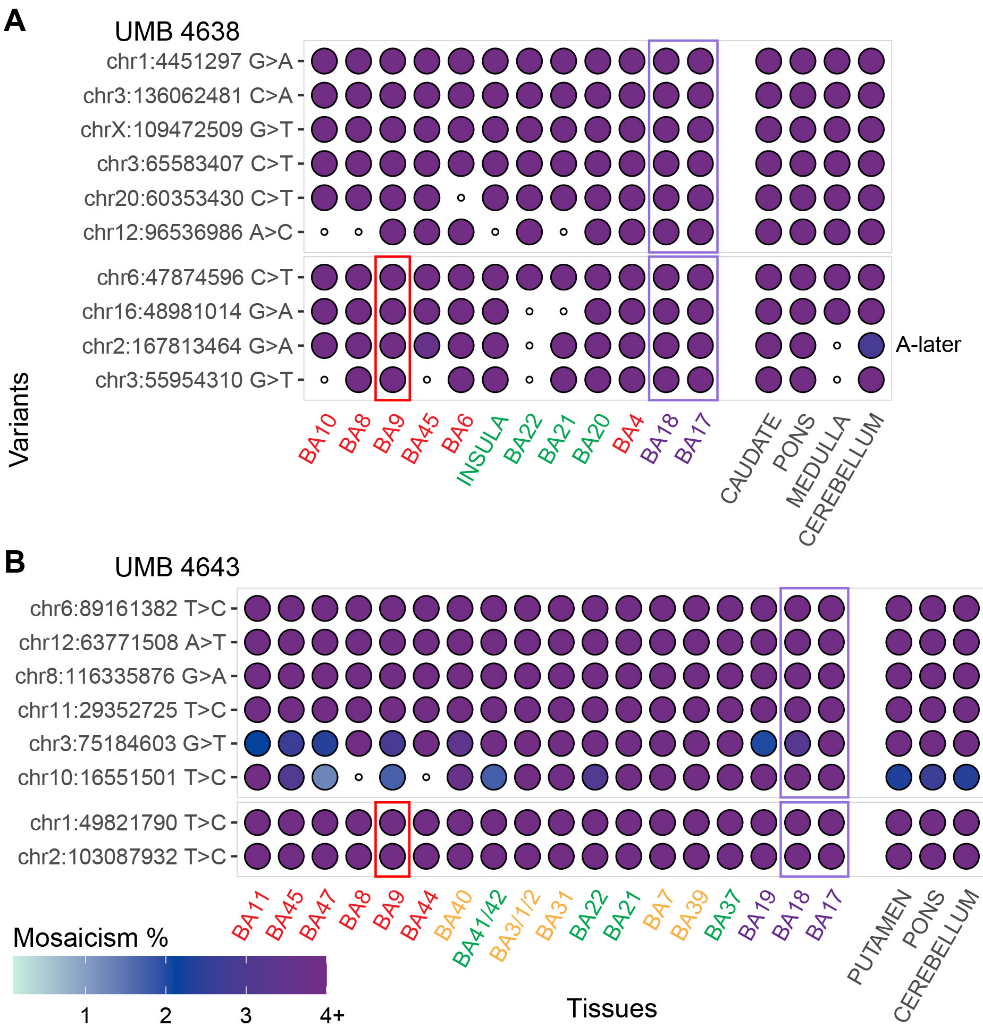


Figure S4: **Mosaicism evaluation of sSNVs originally detected in multiple regions across the left hemisphere, Related to Fig. 3 and Fig. 4.** **(A and B)** Mosaic mutations detected in multiple cortical regions, such as adjacent cortical sites (BA17 and BA18, purple rectangles) or across the cortex (BA9, BA18, and BA17, red and purple rectangles) are often seen shared across almost all sampled cortical tissues as well as most non-cortical samples at similar levels of mosaicism. UMB4638 mutations are represented in **(A)**, and UMB4643 mutations are represented in **(B)**. Regions are indicated by text color: frontal cortex (red), parietal cortex (yellow), temporal cortex (green), occipital cortex (purple), and non-BA regions (black).

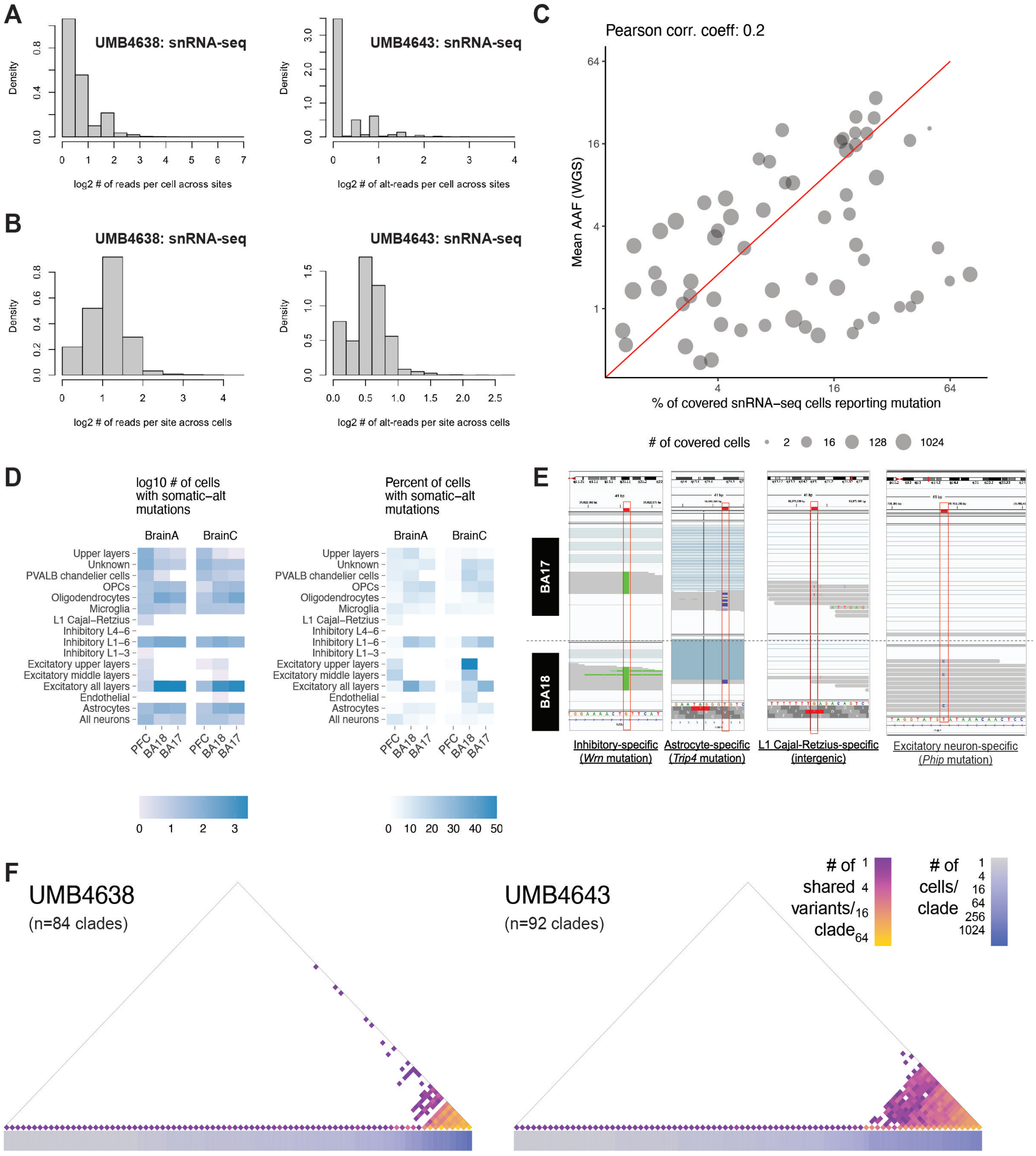


Figure S5: **Coverage statistics and basic properties of sSNVs captured in snRNA-seq and snATAC-seq data. Related to Fig. 5.** **(A)** Log-scaled number of reads per cell summed up across sites that cover the location of a known somatic-alt variant for UMB4638 (left) and UMB4643 (right) in snRNA-seq. **(B)** Log-scaled number of reads per site summed up across cells that cover the location of a known somatic-alt variant for UMB4638 (left) and UMB4643 (right) in snRNA-seq. **(C)** Correlation of the cell fraction of each sSNV (at <50% AAF) in snRNA-seq with the mean AAF from WGS. For each sSNV's cell fraction calculation, only cells that have reads overlapping locus are considered in the denominator. **(D)** Number and percentage of cells per annotated cell type that shows coverage of somatic alternative alleles in the data. **(E)** IgV plots of example variants captured in inhibitory neurons, astrocytes, L1 Cajal-Retzius cells, and excitatory neurons. **(F)** Groups of cells that are found to share mutations in snRNA-seq and snATAC-seq data. Groups ("clades") are arranged along the diagonal of the half-square, with off-diagonal entries colored by the number of variants mutually shared between groups.

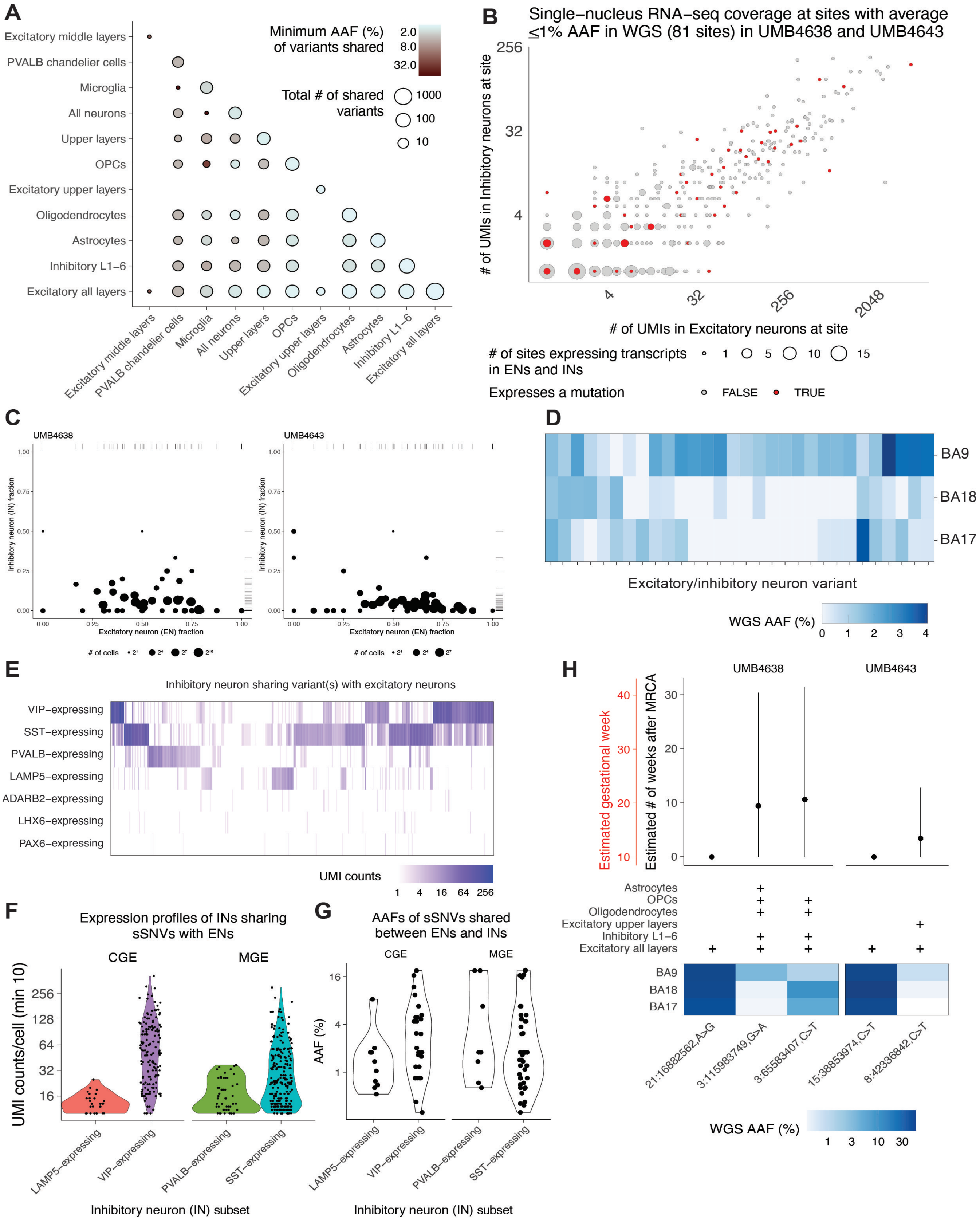


Figure S6: Continued on the next page.

**Transcriptional properties and timing of low-mosaic clones containing excitatory and inhibitory neurons. Related to Fig. 6.** **(A)** The number and estimated minimum AAF of variants shared amongst different cell types. All variants studied across both brains are depicted. **(B)** The number of UMIs (i.e., transcript counts) detected across single excitatory and inhibitory neurons (ENs and INs, respectively) from snRNA-seq at sites with mutant alleles detected at <1% AAF in WGS data. If a site's UMIs support the presence of mutant alleles, points are colored red. Points are sized by the number of sites with the observed pairing of excitatory and inhibitory UMI counts. **(C)** Observed fractions of cells in variant-sharing subgroups that are annotated as excitatory or inhibitory neurons. These raw fractions, along with cell numbers and experimental metadata, were used to generate Empirical Bayes estimates of the cellular composition of each subgroup after controlling for biological and technical factors (**Fig. 6D**). **(D)** Heatmaps of WGS AAFs for mutations detected in both excitatory and inhibitory neurons at <4% WGS across all three cortical regions. **(E)** UMI counts of sets of inhibitory neuron marker transcripts as determined by the Allen Brain Atlas (Hodge et al., 2019). UMI counts are computed for all inhibitory neurons that share at least one variant with an excitatory neuron in the dataset. **(F)** Distributions of marker counts per cell as in **(B)** as grouped by subtypes and region-of-origin of inhibitory neurons. MGE: medial ganglionic eminence. CGE: caudal ganglionic eminence. Only cells with at least 10 UMIs of transcripts corresponding to the marker set are plotted. **(C)** Distributions of AAFs of variants shared between excitatory neurons and inhibitory neurons expressing either CGE or MGE markers. **(H)** Estimated time-of-origins and spatial distributions of variants marking different cell types. Top: Time-of-origin estimates given in PMW (black) or GW (red) for select variants identified in different cell types in UMB4638 (left) and UMB4643 (right). Middle: Cell types from snRNA-seq in which variants were detected are shown. Bottom: Heatmaps showing the WGS AAF estimates for variants in different regions are also shown. The absence of a "+" for a particular cell type only indicates that at least one mutant allele-backing read was detected in that cell type. The absence of a "+" for a cell type does not necessarily mean that only ref reads were detected in the cell type.

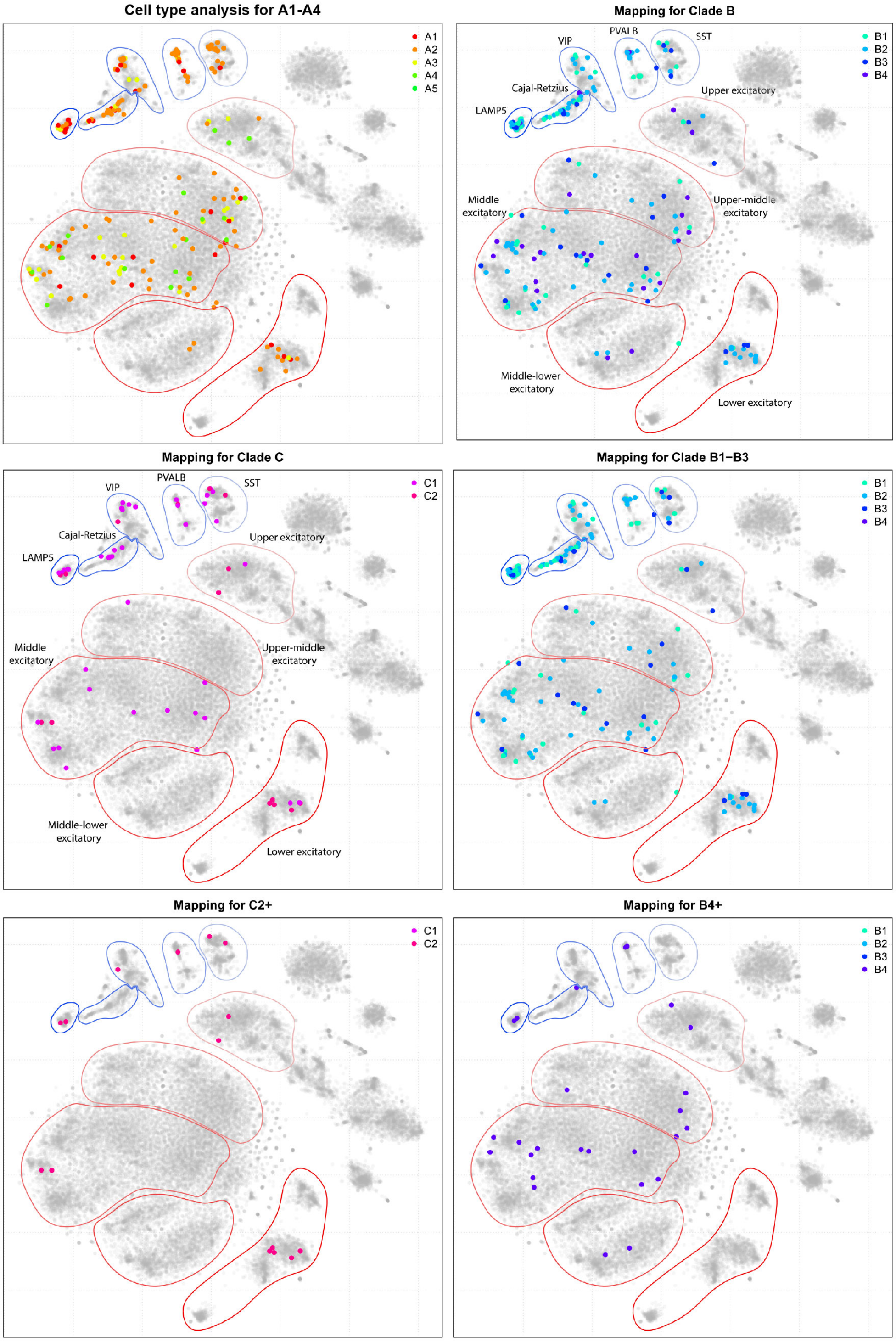


Figure S7: **Cell type analysis of clonally related sSNVs in two additional clades. Related to Fig.7.** PRDD-seq-evaluated sSNVs across subclades A1-4 and two additional clades (B and C) and their corresponding subclades in UMB4638 distribute throughout different excitatory layers and inhibitory subtypes. Neuronal annotations comprise upper, middle, and lower layers of excitatory neurons and four inhibitory interneuron subtypes. PRDD-seq cell type and layer identifications were determined by mapping PRDD-seq cells onto reference 10X Chromium snRNA-seq data performed on three cortical areas (bulk and single cell data sets from BA9, BA18, and BA17). Mapping was conducted based on expression profile similarity of marker genes as described before (Huang et al., 2020) Cells captured and measured with PRDD-seq are mapped on top of clones, which are ordered from earlier occurring (e.g., B1) to later-occurring (B4) within their respective clades.
